# NIPBL and STAG1 enable loop extrusion by providing differential DNA-cohesin affinity

**DOI:** 10.1101/2024.10.22.619583

**Authors:** Raman van Wee, Roi Asor, Yiwen Li, David Drechsel, Maria Popova, Gabriele Litos, Iain Davidson, Jan-Michael Peters, Philipp Kukura

**Affiliations:** Physical and Theoretical Chemistry, Department of Chemistry, Kavli Institute for Nanoscience Discovery, Dorothy Crowfoot Hodgkin Building, University of Oxford, South Parks Road, Oxford OX1 3QU, UK; Department of Physiology, Anatomy and Genetics, University of Oxford, South Parks Road, Oxford, OX1 3QX, UK; Research Institute of Molecular Pathology (IMP), Vienna BioCenter, 1030 Vienna, Austria; Vienna Biocenter Core Facilities, Dr. Bohr-Gasse 7, 1030 Vienna, Austria

## Abstract

DNA loop extrusion by cohesin has emerged as a critical pathway for chromosome organization. In vitro single-molecule experiments indicate that loop extrusion requires the assembly of an heteropentameric complex consisting of the SMC1/SMC3 heterodimer, STAG1, NIPBL and the kleisin SCC1. The multimeric nature of this complex and regulatoin of its dynamic DNA interactions that are modulated by ATP binding-hydrolysis cycles make it challenging to reveal the molecular mechanism of loop extrusion. Here, we use mass photometry to quantify the key interactions responsible for cohesin assembly, DNA binding, and their modulation by ATP binding and hydrolysis. We find that STAG1 binds tightly to the trimeric complex formed by the SMC1/SMC3 heterodimer and SCC1, creating a DNA binding site whose strength is modulated by the presence of ATP. NIPBL binds strongly to both DNA and the STAG1-tetramer, enabling assembly of the holoenzyme in the absence of DNA or ATP. Taken together, these results suggest that NIPBL acts as a DNA anchor, while a dynamic STAG1-Trimer binding site drives DNA loop formation.

## Introduction

Cohesin is responsible for genome organization during interphase^1-4^ enabled by its ability to bind DNA and then reel sequences flanking the binding site into a loop that increases in size over time. This phenomenon is known as loop extrusion and has been visualised directly using single-molecule experiments^5,6^. Cohesin function relies on the formation of a holoenzyme consisting of the SMC1/SMC3 heterodimer, the largely intrinsically disordered kleisin SCC1 (also known as RADS21 or Mcd1), and the HAWKs^7^ STAG1/STAG2/STAG3, and NIPBL (**Fig. 1a**)^2,4^. SMC1 and SMC3 form extended coiled-coil arms that dimerise at a ‘hinge’ interface at one of their ends to adopt a V-shape. The other end of each arm contains an ATPase domain that is related to those found in ATP binding cassette (ABC) transporters^8^, which are connected by SCC1, forming a ring-structure to which STAG1 and NIPBL bind^9^. Mutations in cohesin subunits or its regulators are associated with several developmental diseases, collectively known as cohesinopathies^10-12^.

**Fig 1:**
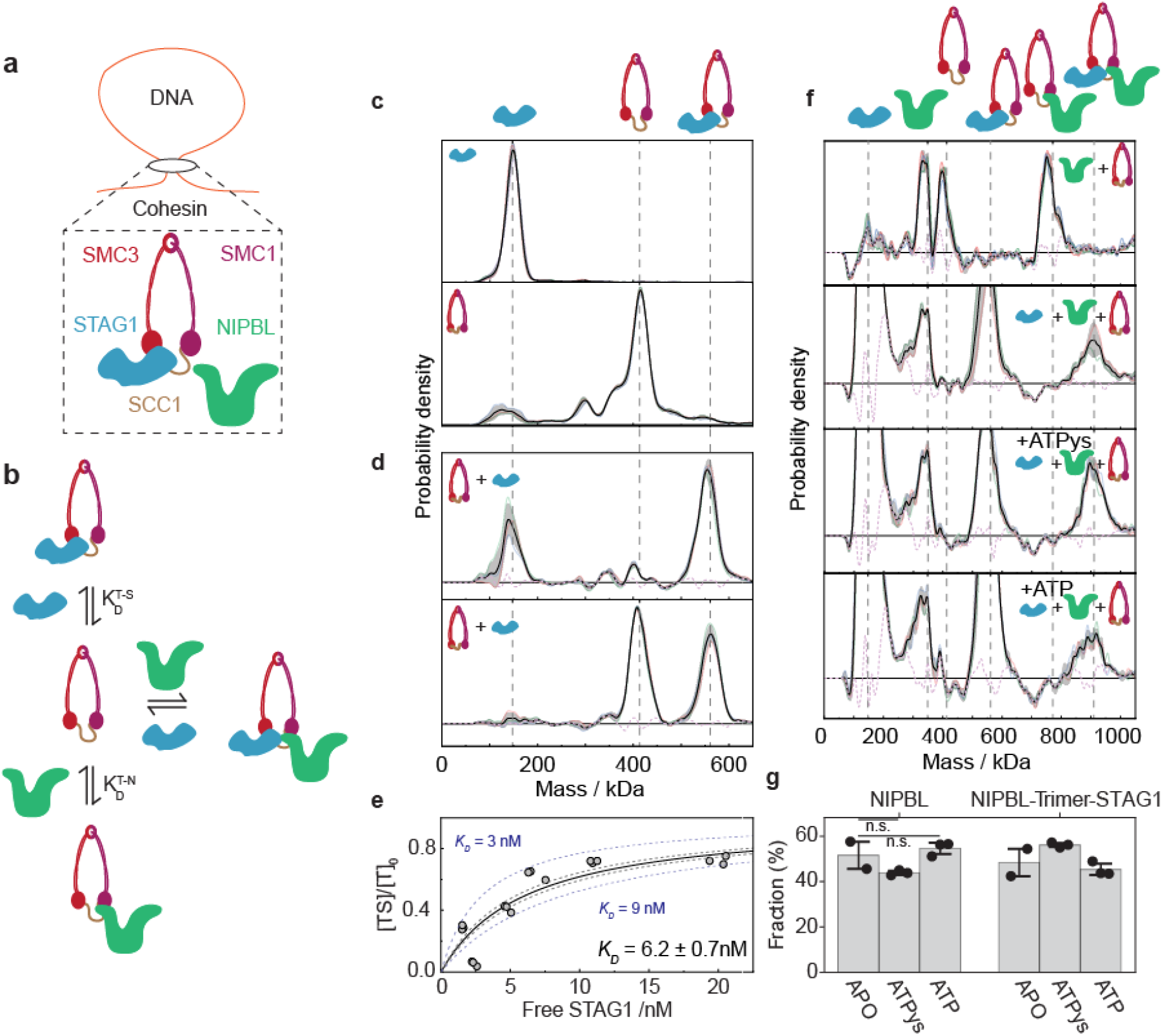
Quantification of the (high) STAG1-Trimer affinity and demonstration of holoenzyme assembly without the need for DNA or ATP (ATP binding or hydrolysis do not affect the stability of the holoenzyme). **a**, Schematic depiction of the cohesin holoenzyme. **b**, Protein – protein interactions involved in the assembly of the holoenzyme. **c**, Representative mass histograms of STAG1 (top) and the Trimer (bottom) with the position of the peak of interest indicated. **d**, Representative mass histograms of Trimer-STAG1 interaction following the mixing of 150 nM STAG1 + 150 nM Trimer (top) and 80 nM STAG1 + 150 nM Trimer (bottom), all total protein concentration. The expected masses of STAG1, Trimer, and STAG1-Trimer are indicated by the vertical dashed grey lines. Both panels show the average (black curve), standard deviation (grey error bars), and between 2 and 3 technical repeats (solid-colored curves). The error of the reconstruction of the experimental histogram from the histograms of the individual components is shown by the dashed purple curve. **e**, Hill plot, showing the molar fraction of the STAG1-Trimer complex as a function of the concentration of free STAG1 (**Fig. S6** for all distributions and fits). All mixtures were incubated at constant Trimer concentration which corresponds to 50 nM total protein concentration (12.6 ± 0.9 nM of effective Trimer concentration, **Fig. S5b**), and a STAG1 concentration ranging from 3 to 30 nM. Circles correspond to six different STAG1 concentrations with 3 technical repeats for each. Solid black curve corresponds to the best fitted Hill equation (see Methods). Gray dashed curves represent the boundaries of the fitting error, and the dashed blue lines correspond to the expected Hill plot for the indicated *K*_D_ values and serve as visual guide for the scale of variation in the estimated dissociation constant. **f**, Mass histograms following the subtraction of the residual signal, emphasizing the detected subunits and complexes of the cohesin holoenzyme (**Fig. S1, Fig. S2**). The subtracted mass histograms correspond to the equilibrated mixture of 150 nM of Trimer sample mixed with 150nM of: NIPBL (top), STAG1 and NIPBL (second from the top), and STAG1 and NIPBL in the presence of 2.5 mM ATP and ATPγS (as indicated). The expected masses of STAG1, NIPBL, Trimer, STAG1-Trimer, NIPBL-Trimer, and STAG1-NIPBL-Trimer are indicated by the vertical dashed grey lines. Each panel shows the average (black curve), standard deviation (grey error bars) of between 2 and 3 technical repeats, indicated by solid-colored curves. The error of the reconstruction of the experimental histogram from the histograms of the individual components is shown by the dashed purple curve. **g**, Molar fractions of free NIPBL and NIPBL that is part of the NIPBL-Trimer-STAG1 complex out of the total NIPBL concentration for the mixtures with STAG1, NIPBL and Trimer (bottom 3 panels in e). Fractions were determined by considering the area under the two peaks. No significant difference is observed for the APO, +ATPγs, or +ATP case (p > 0.05). Bars indicate the mean value across technical repeats (black dots) and black error bars represent the standard deviation.

Structural studies have shown that cohesin adopts a variety of conformations at different stages of the ATPase cycle, including an open ring-shape, a closed rod-shape, and a bent state^13-16^. Following these findings, multiple models for loop extrusion have been proposed, including walking, pumping, scrunching, ratchet and reel and seal models^2,4,17,18^. While parts of these models are supported by structural evidence^13-16,19-26^, the field is yet to agree on a mechanistic description for loop extrusion^2,3,18^.

A key challenge to our understanding of the molecular mechanism underlying cohesin’s loop extrusion function is a lack of quantitative information of the underlying biomolecular interactions. Given that biomolecular mechanisms are controlled by molecular affinities and on-off rates, such information is critical to the development and validation of any proposed mechanism. Structural studies provide atomically-resolved detail, but in the case of cohesin operate at orders of magnitude higher concentrations than those in the cell and are intrinsically dominated by homogeneous, highly abundant complexes that do not necessarily represent the functional species. Single-molecule fluorescence-based studies provide dynamic information but cannot quantify stoichiometries and yield mostly digital on/off information rather than affinities.

Here, we use mass photometry (MP) to quantify the key interactions responsible for cohesin assembly, DNA binding, and their modulation by ATP *in vitro*. Our results show that STAG1 and NIPBL tightly bind the cohesin Trimer, but not each other and that the holoenzyme can assemble in the absence of DNA and ATP. We find that STAG1 and the Trimer each interact weakly with DNA but bind cooperatively to DNA. The affinity of the STAG1-Trimer complex for DNA is reduced in the presence of ATP, while the strong interaction of NIPBL with DNA anchors the holoenzyme on DNA during ATP binding and hydrolysis. Our results are consistent with a model where the STAG1-Trimer complex grabs and releases DNA in an ATP-dependent manner while NIPBL holds on to DNA, thereby leading to loop extrusion.

## Results

### Cohesin assembles without ATP binding or hydrolysis

Our approach to quantifying the underlying affinities is based on mass photometry (MP), which provides a mass-resolved picture of the solution composition of biomolecules and the complexes they form^27,28^. MP measurements of human STAG1 (148 ± 2 kDa, μ ± s.d.) and the SMC1-SMC3-SCC1 Trimer (414 ± 1 kDa, μ ± s.d.) point to varying degrees of polydispersity and purity in native conditions, which are obscured in gel electrophoresis under denaturing conditions (**Fig. 1C, Fig. S1**). To isolate the mass distribution of species participating in the interaction with STAG1, we masked residual impurities in our Trimer preparation from the histogram using a simple spectral decomposition (**SI and Fig. S2**). This subtraction procedure allowed us to visualise changes in relative abundance when using either an excess of Trimer or STAG1 (**Fig. 1d, Fig. S3**). Given the analyte concentrations used, these distributions indicate a low nanomolar affinity, in agreement with the observation that Trimer and STAG1 are frequently co-purified as the ‘cohesin tetramer’^29,30^.

To better quantify the affinity, we performed a titration, ensuring that the mixture was incubated for longer than the time required to equilibrate (**Fig. S4**). A subsequent fit to the Hill equation yields *K*_d_ = 6.2 ± 0.7 nM in close agreement with the value obtained from evaluating individual mass distributions directly^31,32^ of *K*_d_ = 5.4 ± 1.7 nM (**Fig. 1e, Fig. S5, Fig. S6**). These results indicate that a titration as well as single shot measurements with MP yield the desired molecular affinities upon equilibration. Using individual mass distributions for simplicity, NIPBL (343 ± 8 kDa, μ ± s.d.) yields a single shot *K*_d_ of 20 ± 4 nM for interaction with the Trimer (**Fig. 1f**, top panel), in good agreement with ITC measurements of NIPBL and the kleisin subunit^33^. We found NIPBL to self-oligomerize and aggregate to some extent (**Fig. S7**) in agreement with previous reports^15,34^, however our measurements show that the most abundant state of the protein is its monomeric form at our experimental conditions.

Upon mixing all three proteins, the complete holoenzyme (905 kDa) assembled, with no observable difference in the relative abundances between the APO, ATPγs & ATP states (**Fig. 1f** bottom three panels and **Fig. 1g. Fig. S8** shows the unsubtracted distributions). The observation that ATP does not affect the 1:1 interactions of STAG1 and NIPBL with the Trimer (**Fig. S9, Fig. S10**) supports the hypothesis that ATP binding or hydrolysis do not affect the affinities forming the holoenzyme. This observation is in line with the finding that STAG1 can be readily co-purified with ATP binding or hydrolysis deficient mutant forms of cohesin^35-37^. We also did not detect a direct interaction between STAG1 and NIPBL (**Fig. S9, Fig. S10**), suggesting that the strong pairwise interactions with the Trimer drive the assembly of the holoenzyme. We cannot, however, exclude the possibility that STAG1 and NIPBL interact directly when part of the complete pentamer complex^19,20^ with a weak interaction that cannot be detected in solution when the Trimer is not present. Overall, our results show that the holoenzyme is stable without DNA and ATP at low nanomolar concentrations.

### STAG1 and the Trimer bind DNA cooperatively

Loop extrusion requires binding of the holoenzyme to DNA. Electrophoretic mobility shift assays have shown that the holoenzyme contains at least five DNA binding sites with estimated affinities in the high nanomolar to micromolar range, all of which are essential for loop extrusion^15^. However, the contribution of each of these interactions to the recruitment of the cohesin complex to DNA is debated^38^. Some reports ascribe this function primarily to NIPBL^6,19,20,39-41^, while others contradict this^42^. To address this question, we began by measuring the interaction of the Trimer and STAG1 with DNA (**Fig. 2a**). First, we compared binding to circular and linearized DNA constructs with the same sequence and found that purified cohesin tetramer (Trimer-STAG1) preferentially binds to the circular, supercoiled construct (**Fig. S11**). Preferential binding to supercoiled and circular DNA over relaxed and linear DNA has also been observed for condensin and Smc5/6^43-45^, presumably because the energetic penalty for DNA deformation upon DNA binding has already been paid^46^. Since cohesin is able to bind to DNA without entrapping it inside its ring structure ^5,47-49^ it is possible that cohesin preferentially binds circular DNA for the same reasons. However, it is also possible that some spontaneous entrapment of DNA inside cohesin ring’s occurred under our assay conditions^50^ and that this contributed to the preferential binding of circular over linear DNA. We therefore used the circular construct in all subsequent experiments.

**Fig 2:**
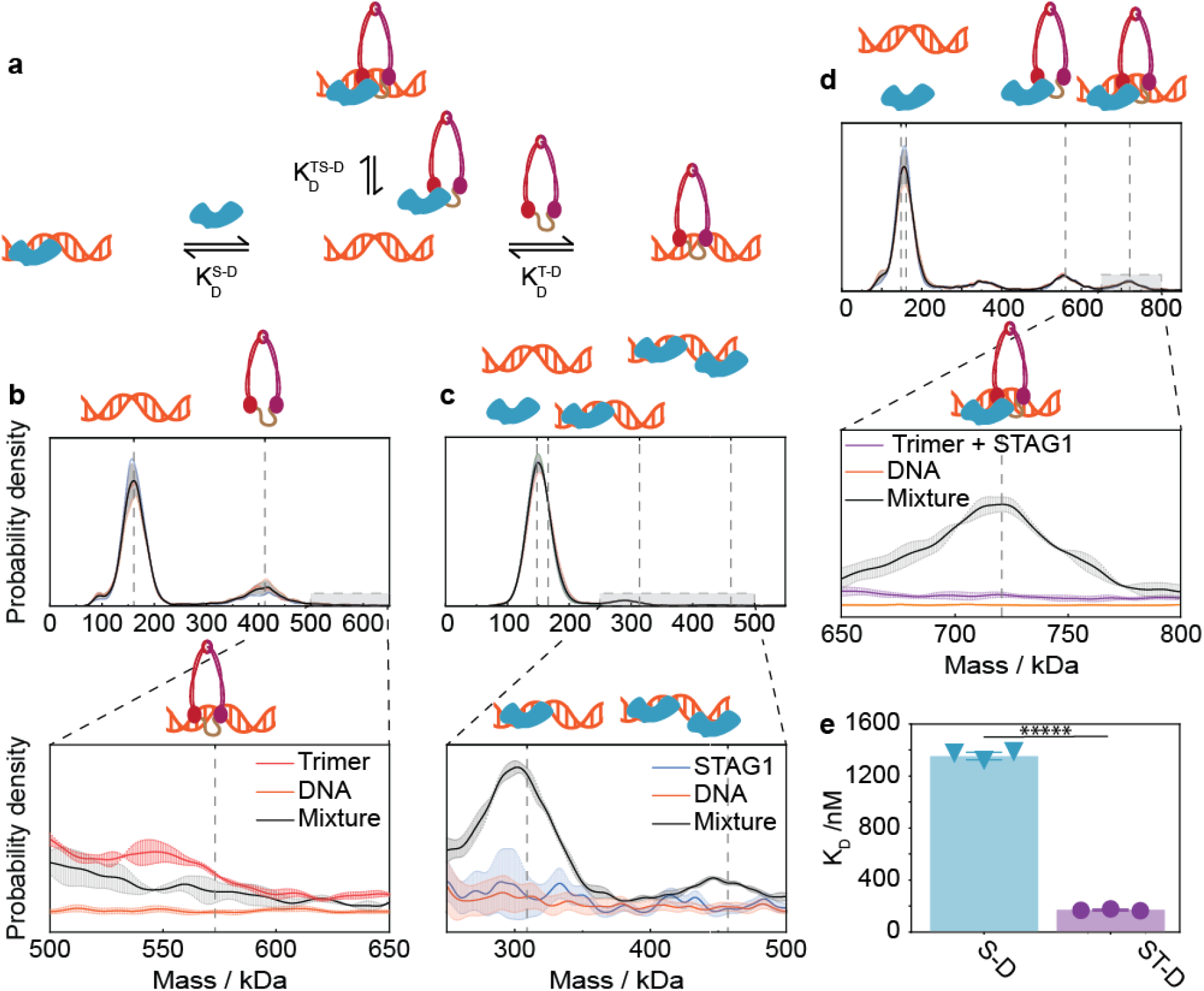
Cooperative interaction of STAG1 –Trimer with DNA. **a**, Illustration of possible interactions of STAG1, Trimer and DNA with their dissociation constants indicated. **b – d**, Top: Mass histograms of mixtures of 150 nM 302 bp minicircular dsDNA with 150 nM Trimer (b), 150 nM STAG1 (c), and 150 nM of Trimer and STAG1 (d). Black curves and grey error bars correspond to the average mass histogram of 3 technical replicates and their standard deviations. Individual mass histograms of the individual replicates are presented as colored curves. For all mass histograms the expected masses of the relevant subunits (DNA, Trimer and STAG1) and their complexes are indicated by the vertical dashed grey lines. Bottom: Mass histograms zoomed around the expected mass of the DNA-protein complexes. For all cases, black curves and error bars correspond to the measured mean distribution and standard deviation of mixtures (b: DNA + Trimer, c: STAG1 + DNA, and d: Trimer + STAG1 + DNA). Colored curves and error bars correspond to the measured mass distribution of the individual components when measured alone. The ratio between the area beneath the mixtures curves (black) and the individual component curves (colored) represent our ability to detect and quantify the formation of the DNA-protein complex (0.54 ± 0.03 (a), 2.73 ± 0.11 and 3.08 ± 0.04 (b), and 11.2 ± 0.02 (c)). Grey dashed vertical lines indicate the expected mass of the Trimer-DNA complex (b), STAG1-DNA 1:1 and 2:1 complexes (c), and STAG1-Trimer-DNA complex (d). **e**, Single-shot *K*_D_ values for the interactions of STAG1 and STAG1-Trimer with DNA, which are 1.35 ± 0.03 µM & 169 ± 6 nM, respectively. Bars indicate the mean value across technical repeats (black dots) and black error bars represent the standard deviation. The *K*_D_ of the Trimer-DNA interaction is beyond the detection limit of mass photometry. Assuming a conservative detection limit of 1 nM, we estimated the lower bound for the *K*_D_ to be >2 µM. Since the STAG1 and DNA are unresolvable, the total area under their mutual peak in (c) was assumed to contain 50% STAG1 and 50% DNA, variations in the extracted *K*_D_ values assuming different ratios are shown in **Fig. S12**. *****: p<0.000005.

Upon mixing DNA and Trimer at equimolar concentrations of 150 nM we observed clear peaks at the expected masses of the Trimer and DNA (164 ± 4 and 414 ± 1 kDa, both mean ± s.d.), but not at the mass of a 1:1 complex (578 kDa) (**Fig. 2b**, top). When zoomed in on the mass range of the Trimer-DNA complex, we found that the signal was indistinguishable to that of the DNA or Trimer when measured individually (**Fig. 2b**, bottom). We therefore conclude that the interaction of the Trimer with DNA is too weak to be observed at the concentrations used and must be in the micromolar range or higher, in line with previous results based on fluorescence polarization^15,51^.

Given that the affinity of the Trimer for DNA is too low to load onto DNA at cellular concentrations^52^, either STAG1, NIPBL, or both must have a facilitating role. STAG1 is known to contain three DNA binding patches^15,51^, and indeed when we incubated STAG1 with DNA, we observed a 1:1 or a 2:1 (STAG1:DNA) interaction (**Fig. 2c**). To test whether STAG1 could contribute to recruiting the Trimer to DNA or retaining it there, we equilibrated STAG1 and Trimer at equimolar concentrations of 150 nM, and then added DNA. We found that when the Trimer is in complex with STAG1, it bound to DNA (**Fig. 2d**). These results add to the body of evidence showing that NIPBL is not strictly required for binding cohesin to DNA *in vitro*^53,54^, although it is frequently referred to as ‘the cohesin loader’^19,20,23,33,39,40,42,54^. Quantification of the STAG1 fraction bound to DNA (**Fig. 2c**) relative to the fraction of STAG1-Trimer on DNA (**Fig. 2d**) reveals that the latter is larger, suggesting a stronger interaction. Indeed, when we quantify the *K*_d_ from these measurements, we find 1.35 ± 0.03 µM for STAG1-DNA and 169 ± 6 nM for the STAG1-Trimer complex (**Fig. 2E, Fig. S9, Fig. S10, Fig. S12**). The 8-fold increase in affinity indicates a cooperative effect, presumably caused by simultaneous interaction of both STAG1 and the Trimer with DNA.

### ATP presence modulates the affinity of the Trimer for DNA

In order to progressively extrude a loop, adjacent segments of DNA must be reeled in during subsequent mechanochemical cycles. Regardless of the molecular mechanism by which cohesin extrudes DNA, the interaction of the complex with DNA must be dynamic to allow a ‘DNA-grabbing’ state with high affinity for DNA and a ‘DNA-release’ state when the newly reeled DNA is released into the loop. Given that cohesin requires ATP to perform loop extrusion^5,6^, we repeated our DNA binding experiments with the STAG1-Trimer complex in the presence of ATPγs (to probe binding) or ATP (to probe hydrolysis). We found that the presence of either nucleotide reduced the STAG1-Trimer affinity for DNA below the detection limit in our assay (**Fig. 3a-b, Fig. S13**). To exclude a buffer dependent effect, we repeated this experiment but replaced the phosphate buffer with a Tris buffer, which resulted in the same observation (**Fig. S14** and **Fig. S15**). Since STAG1 does not have an ATP binding domain and is not released from DNA in the presence of these nucleotides when measured alone (**Fig. S9 & Fig. S10**), we hypothesize that the affinity of the STAG1-Trimer for DNA is reduced upon ATP binding at one (or both) of the Trimer ATPases, due to their proximity to DNA^20^.

**Fig 3:**
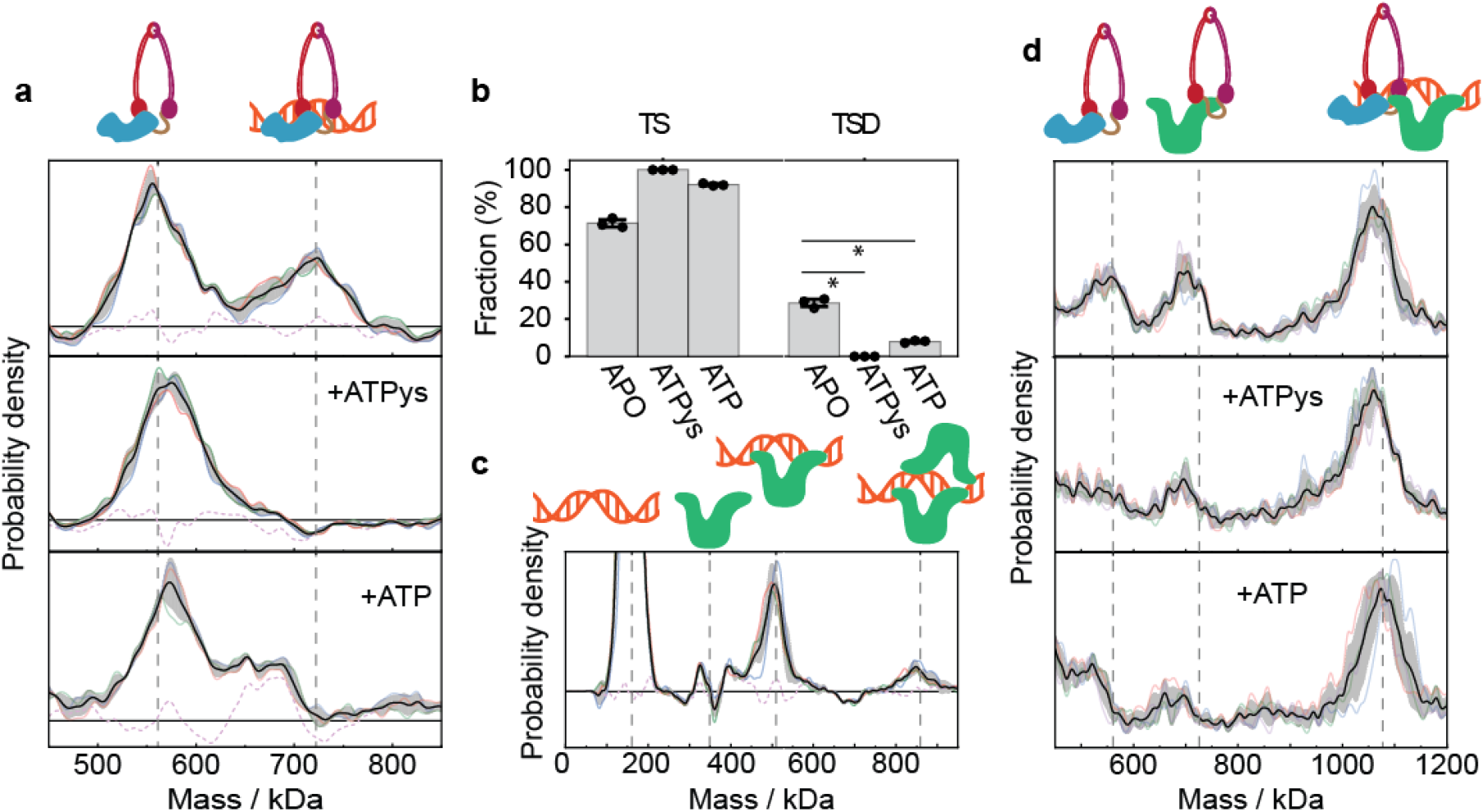
The STAG1-Trimer complex is released upon ATP binding while NIPBL stabilizes the full pentameric complex on DNA. **a**, Mass distributions with residual spectrum subtracted for equilibrated mixtures of STAG1, Trimer and DNA all present at 150 nM (top panel), and the same mixture in the presence of 2.5 mM ATPγs (middle panel) or 2.5 mM ATP (bottom panel). Each panel shows the expected peak location of STAG1-Trimer and STAG1-Trimer-DNA complexes (vertical dashed grey lines). Black curves and grey error bars correspond to the averages and standard deviations of 3 technical repeats per experiment, respectively. Colored curves correspond to the individual repeats and dashed purple curve is the error of reconstructing the mass distribution from the individual components (**Fig. S2**). **b**, The abundance of STAG1-Trimer and STAG1-Trimer-DNA relative to the total STAG1-Trimer abundance for the mixtures in (a) as determined by the area of the Gaussian fit to the peaks of interest, showing a significant difference for the APO, +ATPγs, or +ATP case (p < 0.05). Bars indicate the mean value across technical repeats (black dots) and black error bars represent the standard deviation. **c**, Residual mass distribution for a mixture of 150 nM NIPBL with 150 nM 302 bp minicircle dsDNA. Average distribution, standard deviation and individual repeats are indicated as in (a). Vertical lines indicating the expected masses of the different molecular species, showing 1:1 and 2:1 binding of NIPBL to DNA. **d**, Mass distribution for mixing STAG1, Trimer, NIPBL and 302 bp minicircle dsDNA all at 150nM. The affinity of NIPBL for the Trimer or STAG1 is not significantly affected by the presence of ATPγs or ATP (**Fig. S9**).

Most interactions of cohesin with DNA are mediated by electrostatic forces with the phosphate backbone^19,20^, hence binding of negatively charged ATP could be a driving force for the reduced affinity. Thus, we examined the effect of pure ionic strength on the STAG1-Trimer-DNA interaction. We incubated the STAG1-Trimer complex with DNA in a solution with the same ionic strength as the solution containing ATP or ATPγs but replaced the nucleotide with NaCl with an equivalent ionic contribution (Materials and Methods). Under these conditions, the STAG1-Trimer did not interact with DNA, whilst the protein complex remained intact (**Fig. S14, Fig. S15**). This behaviour is consistent with the recent observation that the abundance of STAG1 on DNA decreased in living cells upon increasing the ionic strength of the incubation buffer^42^ and suggest that indeed electrostatic interactions are crucial for STAG1-Trimer binding to DNA. Although we cannot distinguish a direct effect due to ATP binding from an indirect effect due to the increased ionic strength, our results show that reduced electrostatic interactions substantially weaken the STAG1-Trimer to DNA affinity, resulting in DNA release. Additionally, we cannot rule out that the affinity is also modulated through structural rearrangements induced by ATP presence, such as head engagement^15,19,55,56^ and kleisin gate opening.

For cohesin to processively extrude DNA, it must stay bound to DNA throughout its mechanochemical cycle. Since the STAG1-Trimer did not detectably bind to DNA in the presence of ATP, this suggests that the DNA binding site(s) on the STAG1-Trimer complex are insufficient. We therefore hypothesized that NIPBL might fulfil this role. Mixing of equimolar concentrations of NIPBL and DNA (150 nM each) revealed peaks at 508 kDa and 851 kDa, indicating direct 1:1 and 2:1 NIPBL:DNA interactions (**Fig 3c** and **Fig. S16**), as for STAG1-DNA (**Fig. 2c**). Since we did not detect free NIPBL in solution, we estimate the upper limit for the dissociation constant to be 10 nM. Our results thus suggest that the NIPBL-DNA interaction is substantially stronger than the micromolar affinity previously reported^15,57^. However, in those studies, N-terminal truncated NIPBL (ΔN-NIPBL) was used. Indeed, when we probed the interaction of ΔN-NIPBL with DNA, we could not detect any complex formation (**Fig. S17**), indicative of a dissociation constant in the micromolar range or higher^15^.

When we mixed the three components of the holoenzyme (STAG1, Trimer and NIPBL) and then added DNA to the mixture, the intact holoenzyme associated with DNA (1069 kDa, **Fig. 3d** and **Fig. S18**), supporting the notion that ATP is not required for DNA binding *in vitro*^34,53,54,58^. Moreover, we did not detect any unbound holoenzyme, as was detected for the interaction between STAG1-Trimer complex and DNA (**Fig. 2d**), showing that the strong interaction of NIPBL with DNA increases the affinity of the holoenzyme for DNA. To assess whether NIPBL might be responsible for retaining cohesin on DNA during a mechanochemical cycle, we repeated this experiment in the presence of ATPγs and ATP. We did not observe release of the cohesin complex, confirming that NIPBL stabilizes the cohesin complex on DNA in the presence of ATP, irrespective of the mechanism of affinity modulation of the STAG1-Trimer-DNA interaction.

## Discussion

Our results show that the affinities of the cohesin Trimer, with both STAG1 and NIPBL are in the low nanomolar range and independent of ATP, while a pairwise interaction between STAG1 and NIPBL was not observed. These results suggest strong and independent interactions of STAG1 and NIPBL with the Trimer. Specifically, the high STAG1-Trimer affinity suggests that the proteins form a tight complex with a slow unbinding rate, which agrees well with previous observations that the two proteins coelute during purification^29,30^. The interaction of the Trimer with NIPBL is ∼3 times weaker than STAG1 and is therefore expected to be more dynamic^5,57,59^. While our results suggest micromolar or higher interaction strength between STAG1 and NIPBL, we cannot rule out an additional STAG1-NIPBL interaction that takes place as part of the pentameric complex of the holoenzyme^20^.

In contrast with the pairwise protein-protein interactions, the multivalent interaction landscape between cohesin subunits and DNA is more complex. These binding sites cooperatively interact with DNA with an apparent affinity that is modulated by ATP or changes in ionic strength. Our results suggest distinct roles for STAG1 and NIPBL in facilitating DNA-protein interactions for cohesin’s mechanochemical cycle (**Fig. 4a-b**). Our finding of a cooperatively-enhanced STAG1-Trimer-DNA interaction (**Fig. 4a**) suggests that STAG1 serves as an ‘DNA affinity enhancer’ for the Trimer, such that the latter can grab, and hold on to DNA. This hypothesis is supported by the fact that STAG1 is not required for cohesin’s ATPase activity^5,37^, does not affect the extent of DNA supercoiling induced by loop extrusion^60^, and is required for loop extrusion^5^ in high salt buffer only^57^. Our results further suggest that the STAG1-Trimer complex is in principle able to bind DNA, but not in the presence of ATP or physiological salt concentrations. The weakening of the STAG1-Trimer-DNA interaction upon ATP addition suggests that under the conditions required for DNA extrusion, STAG1 cannot establish a sufficiently strong connection to DNA to serve as an ‘anchor’ in the extrusion process.

**Fig 4:**
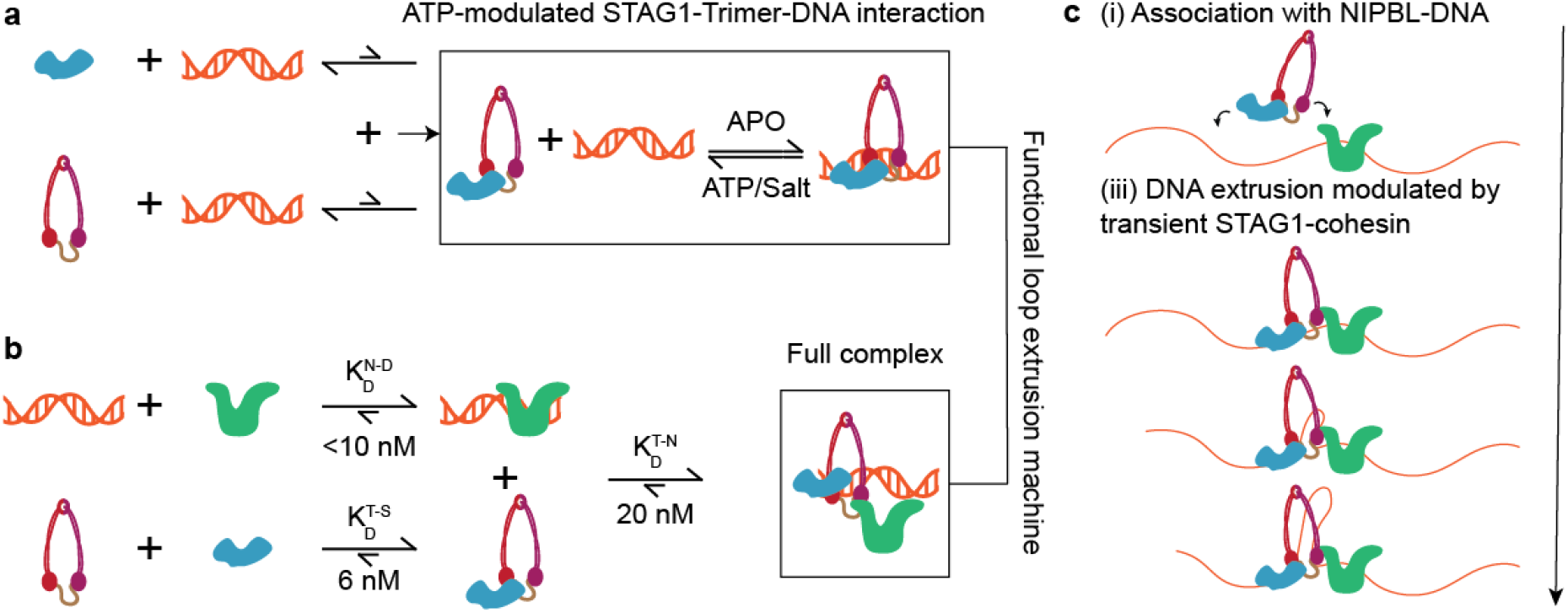
Quantitative description of the fundamental interactions governing cohesin-DNA affinity and its implication on loop extrusion. Schematic overview of the biomolecular interactions characterized in this work. The relative lengths of arrows indicate the equilibrium of the reaction. **a, b**, Interaction strengths resulting in assembly of an ATP-modulated Trimer-STAG1-DNA interaction and the full complex. **c**, Proposed model for loop extrusion, based on characterization of the interaction strengths in (a) and (b) and the existing literature.

Out of the three tested holoenzyme subunits, NIPBL exhibited the highest affinity to DNA. While the interaction of NIPBL and the Trimer (20 nM) is weaker than STAG1, it is sufficient to facilitate assembly of the complete holoenzyme on DNA. Our results suggest that in addition to its effect on cohesin’s ATPase activity^5,35,37^, NIPBL is not strictly required for binding of cohesin to DNA, but stabilises the complete complex on DNA, in line with previous work that observed stabilization on chromatin^42^, especially in the presence of ATP. It is important to note that high affinity and a dynamic nature of the interaction are not mutually exclusive. Since the 1:1 interaction between NIPBL and DNA is not sequence specific and electrostatic in nature^61,62^, long-ranged electrostatic attractions are expected to result in a relatively fast association rate. If we assume that the rate is on the higher reported end for electrostatic interactions (10^8^ − 10^9^ *M*^−1^*s*^−1^)^63^, the dissociation rate constant would be 0.1 – 1 *s*^−1^. This corresponds to an average life time of 1 - 10 seconds given the upper limit of the dissociation constant of 10 nM for NIPBL-DNA interaction determined in this work. We expect the lifetime of the complete complex to be longer, as multiple interactions are involved in the stabilisation of the pentameric complex with DNA. Given typical loop extrusion rates of 0.5 – 2 kb/s^5,6^ and typical loop sizes of 33 kb ^6^, the expected lifetime of the functional complex is tens of seconds, which matches our calculation above and the experimentally observed lifetime of ∼50 seconds for NIPBL and cohesin-NIPBL^5,59^.

Our quantitative description of the interactions of the cohesin holoenzyme reveal that the binding of both STAG1 and NIPBL enable the Trimer to interact with DNA. However, the two proteins provide differential affinity. STAG1 interacts strongly with the Trimer and provides moderate affinity to DNA that can be readily modulated by the presence of ATP or ionic strength. NIPBL on the other hand binds the Trimer more weakly but provides a high affinity to DNA that is independent of ATP. Overall, our results are consistent with the following model for loop extrusion (**Fig. 4c**). Given the concentrations of STAG1 (70 nM), the Trimer (330 nM for SCC1), and NIPBL (200 nM) inside the nucleus^52^, we predict that STAG1 and the Trimer exist as a complex, while a subpopulation of NIPBL associates with DNA and another interacts with the STAG1-Trimer complex. The STAG1-Trimer complex is stabilised on DNA by binding naked DNA and a second interaction mediated by NIPBL. The complete holoenzyme binds DNA more tightly than any of the individual proteins, explaining why the chromosome occupancy by NIPBL is significantly reduced upon depleting cohesin^59^. The dynamic ATPase head on the Trimer reels in DNA by undergoing consecutive ATP-dependent structural changes and affinity modulations, while NIPBL remains bound to DNA. Owing to the proximity to the DNA, a second weak DNA binding site on the Trimer prevents newly extruded DNA from slipping.

This ‘anchor’ function of NIPBL agrees with the observation that PDS5, which is a binding partner of the cohesin release factor WAPL, competes with NIPBL for binding to the kleisin^33,37,64-67^. Moreover, the hypothesis that the STAG1-Trimer predominantly interacts with NIPBL on DNA, and not in solution, is consistent with data in Xenopus showing that most soluble cohesin complexes do not contain NIPBL^68,69^ and that some chromatin have been reported to remodelers recruit NIPBL to chromatin^70^. Furthermore, since it is known that CTCF barriers do not get extruded and instead accumulate at the base of the loop, it seems plausible that CTCF acts on the dynamic part of the extruder, which it encounters first. In our model this would be STAG1, and indeed, a direct interaction between STAG2 and CTCF has been shown to be critical for the location of loop extrusion^71^.

Iit might appear the functional assignments of HEAT II (STAG1) and HEAT I (NIPBL) in this model are swapped relative to what has been proposed for condensin in the hold-and-feed model^24^. However, this work indicated that the anchor role is performed by a topological chamber formed by HEAT II and the Trimer, and not by HEAT II itself. Our observation and those of Shaltiel *et al*. are therefore not mutually exclusive. Moreover, the ΔHEAT II mutant did not have a reduced lifetime on DNA, which could be explained by our model in which HEAT II dynamically grabs DNA, but HEAT I performs the anchoring role. We note that our model is compatible with the Brownian Ratchet models of cohesin mediated DNA loop extrusion^72^. Further work will be required to understand how the DNA binding affinities of each subunit are affected during ongoing loop extrusion.

In summary, we have used MP to quantify all the biomolecular affinities in the cohesin holoenzyme, thereby gaining detailed, mechanistic information. Our data reveal a complex interaction landscape where affinities span at least 3 orders of magnitude in strength and are differentially modulated by ATP in a way that is key to the overall function of the cohesin machine. Our study suggests that the increased complexity introduced by incorporating NIPBL and STAG1 helper proteins allows the modulation of local binding affinities that further facilitate the functionality of the cohesin complex. We envision similar investigations on related SMCs will elucidate mechanistic similarities and differences within this protein family. The development of assays to directly quantify the molecular composition of a loop extruding cohesin complex will test our proposed model, whilst providing rich insights about the interaction dynamics.

## Acknowledgements

We thank the staff at IMBA/IMP/GMI Molecular Biology Service for technical support and Peters lab members for discussions. J-MP is also an adjunct professor at the Medical University of Vienna.

## Funding

Wellcome Trust, Grant Number: 218514/Z/19/Z (RW)

EMBO long-term postdoctoral fellowship ALTF-198-2020 (RA) Boehringer Ingelheim (J-MP)

Austrian Life Sciences Programme 2023 (LS23 IF, project FO999902549) (J-MP)

European Research Council Horizon 2020 Research and Innovation Programme 101020558 (J-MP) Human Frontier Science Program RGP0057/2018 (J-MP)

Vienna Science and Technology Fund LS19-029 (J-MP)

European Research Council PHOTOMASS 819593 (PK)

Engineering and Physical Sciences Research Council EP/T03419X/1 (PK)

## Open Access

This research was funded in whole or in part by ERC (PHOTOMASS 819593) EPSRC EP/T03419X/1. For the purpose of Open Access, the author has applied a CC BY public copyright licence to any Author Accepted Manuscript (AAM) version arising from this submission.

