## Supplementary Information for "NIPBL and STAG1 enable loop extrusion by providing differential DNA-cohesin affinity"

**This PDF file includes:**
**Materials & Methods section**
**Figures S1 to S17**
**Tables S1 to S2**
**SI References**

### Materials and Methods

| Reagent or resource | Source | Identifier |
|---|---|---|
| Adenosine 5′-triphosphate disodium salt hydrate | Sigma-Aldrich | A26209 |
| Adenosine-5'-(γ-thio)-triphosphate, Tetralithium salt | Jena Bioscience | NU-406 |
| Magnesium Chloride hexahydrate | Sigma Aldrich | M2670 |
| HEPES | Sigma Aldrich | H3375 |
| Trizma base | Sigma Aldrich | T1503 |
| NaCl | Sigma Aldrich | S3014 |
| NaH2PO4 | Acros Organics | 3898725000 |
| NaOH | VWR | 28244.262 |
| Glycerol | Sigma Aldrich | G5516 |
| (3-Aminopropyl)triethoxysilane (APTES) | Sigma Aldrich | 440140 |
| Acetone | Sigma Aldrich | 34850-M |
| Glass cover slip | VWR | 630-2867 |
| CultureWell gaskets | Merck | GBL103250 |
| AcquireMP | Refeyn Ltd. | AcquireMP |
| DiscoverMP | Refeyn Ltd. | DiscoverMP |
| Sf900 III SFM cell culture medium | Thermo Fisher | 12658019 |
| Fugene 6 transfection reagent | Promega | E2691 |
| Gibco Fetal bovine serum | Fisher Scientific | 10-082-147 |
| Penicillin-Streptomycin | Sigma Aldrich | P4333 |
| Gibco L-glutamine | Fisher Scientific | 25-030-081 |
| Toyopearl AF-chelate-650M resin | Tosoh Bioscience | 0014475 |
| Strep-Tactin® Sepharose® resin | IBA Lifesciences | 2-1201-002 |
| cOmplete EDTA-free protease inhibitor cocktail | Sigma Aldrich | COEDTAF-RO |

| | | |
|---|---|---|
| Vivaspin 20 100kDa molecular weigh cut off (MWCO) spin filter | Sigma Aldrich | GE28-9323-63 |
| L-arabinose | Sigma Aldrich | A3256 |
| E. coli BL21 | New England Biolabs | C2530H |
| kanamycin | Sigma Aldrich | K4000-5g |
| NucleoBond Xtra Maxi kit | Macherey-Nagel | 740414.50 |
| PEG6000 | FLUKA | 81260 |
| T5 exonuclease | New England Biolabs | M0663L |
| BamHI-HF | New England Biolabs | R3136L |
| HindIII-HF | New England Biolabs | R3104L |
| Monarch Clean-up kit | New England Biolabs | T1030 |
| Ethylenediaminetetraacetic acid (EDTA) | PanreacApplichem | A1103 |
| PstI digestion enzyme | Thermo Fisher | FD0614 |

#### Protein expression and purification

pLib, pBig1a, pBig1b and pBig2ab vectors were transformed into DH10EmBacY cells and bacmid DNA was isolated as described.[1] Spodoptera frugiperda Sf9 insect cells were seeded on a 6 well plate in Sf900 III SFM (Thermo Fisher) and transfected with bacmid DNA using Fugene 6 (Promega). The baculovirus-containing cell supernatant (V0) was collected 96 hours after transfection and used to infect 50 ml Sf9 cells at a density of 1 x$10^6$ cells/ml. Cells were grown at 100 rpm and 27 °C in Grace medium supplemented with Fetal bovine serum (Gibco), Penicillin-Streptomycin (Sigma), L-glutamine (Gibco) and Poloxamer 188 (Gibco) and centrifuged 72 hours after infection. The cell supernatant (V1) was then used to directly infect expression cultures at a density of 1.2 x$10^6$ cells/ml. Expression cultures were grown in Grace medium at 100 rpm and 27 °C. Cells were centrifuged around 54 hours after infection, washed in PBS, frozen in liquid nitrogen and stored at -80 °C.

Tetrameric wild type cohesin was expressed either by co-infecting cells with two baculoviruses generated from pBig1a SMC1/SMC3-Flag (#c190) and pBig1b SCC1-Halo/10xHis-STAG1 (#c350) or by infecting cells with a single baculovirus generated from pBig2ab SMC1/SMC3-Flag/SCC1-Halo/10xHis-STAG1 (#c357). KA/KA cohesin was generated by co-infecting cells with two baculoviruses generated from pBig1a SMC1$^{K38A}$ /SMC3$^{K38A}$-Flag (#c130) and pBig1b SCC1-Halo/10xHis-STAG1 (#c350). All tetrameric cohesin complexes contained the mutations R172A/D279A/R450A in SCC1 that prevent caspase and separase cleavage.[2]

All protein purifications were performed at 4 °C. Tetrameric cohesin, NIPBL (pLib TwinStrep-Flag-NIPBL-10xHis; #712C or pLib Flag-Halo-NIPBL-10xHis; #c253) and STAG1 (pLib 10xHis-STAG1; #c179) were purified as described[3].
ΔN-NIPBL (pLib TwinStrep-Flag-Halo-NIPBL (Δ22-1040)-10xHis; #BB17/241) was purified as described[4] with modifications. Cells were thawed and resuspended in NIPBL purification buffer 1 (50 mM $NaH_2PO_4/Na_2HPO_4$ pH 7.6, 500 mM NaCl, 5 % glycerol, 0.1 % Tween-20) supplemented with 10 mM imidazole pH 7.5, 1 mM PMSF, 1 mM benzamidine, 3 mM betamercaptoethanol and cOmplete EDTA-free protease inhibitor cocktail (Sigma) and lysed by Dounce homogenization. After centrifugation (48000 g, 45 min), the soluble fraction was combined with 5 ml of Toyopearl AF-chelate-650M resin (Tosoh Bioscience) pre-charged with $Ni^{2+}$ ions and incubated for 3 h. The resin was washed with 3 x 10 resin volumes of NIPBL purification buffer 1 supplemented with 15 mM imidazole pH 7.5. Bound protein was eluted with 25 ml NIPBL purification buffer 2 (50 mM $NaH_2PO_4/Na_2HPO_4$ pH 7.5, 150 mM NaCl, 300 mM imizazole pH 7.5, 5 % glycerol. The eluate was mixed with 4 ml of Strep-Tactin Sepharose (IBA Lifesciences 2-1201-025) and incubated for 3 hours. The resin was washed with 3 x 10 resin volumes of NIPBL purification buffer 3 (50 mM $NaH_2PO4/Na_2HPO_4$ pH 7.6, 100 mM NaCl, 50 mM imidazole pH 7.5, 5 % glycerol) and eluted with 20 ml NIPBL purification buffer 3 supplemented with 8mM of desthiobiotin. The eluate was concentrated using a Vivaspin 20 100kDa molecular weight cut off (MWCO) spin filter (Sigma), frozen in liquid nitrogen and stored at -80 °C.

Trimeric cohesin (pBig2ab SMC1, SMC3-Flag, SCC1(TEV)-Halo-14xHis; #BB26) expressing cells (~ 12 ml cell pellet) were thawed and resuspended in cohesin purification buffer 1 (50 mM $NaH_2PO_4/Na_2HPO_4$ pH 7.5, 500 mM NaCl, 5 % glycerol) supplemented with 10 mM imidazole pH 7.5, 1 mM PMSF, and cOmplete EDTA-free protease inhibitor cocktail (Sigma) and lysed by Dounce homogenization. After centrifugation (48000 g, 45 min), the soluble fraction was combined with 5 ml of Toyopearl AF-chelate-650M resin (Tosoh Bioscience) pre-charged with $Ni^{2+}$ ions and incubated for 3 h. The resin was washed with 3 x 10 resin volumes of cohesin purification buffer 1 supplemented with 15 mM imidazole pH 7.5. Bound protein was eluted with 25 ml cohesin purification buffer 2 (50 mM $NaH_2PO_4/Na_2HPO_4$ pH 7.5, 150 mM NaCl, 300 mM imizazole pH 7.5, 5 % glycerol). The eluate was then combined with 5 ml of FLAG-M2 agarose resin (Sigma; A2220) and incubated for 2 hours. The resin was washed with 3 x 10 resin volumes of cohesin purification buffer 3 (50 mM $NaH_2PO_4/Na_2HPO_4$pH 7.5, 150 mM NaCl, 5 % glycerol, 50 mM imidazole pH 7.5). Bound protein was eluted with 25 ml cohesin purification buffer 3 supplemented with 0.25 mg/ml 3xFlag peptide and applied to 0.5 ml Poros HS resin pre-washed with H2O and cohesin purification buffer 3 and incubated for 30 minutes. The resin was transferred to a spin column and eluted four times with 150 µl cohesin purification buffer 4 (50 mM $NaH_2PO_4/Na_2HPO_4$ pH 7.5, 750 mM NaCl, 5 % glycerol, 50 mM imidazole pH 7.5). Eluates were pooled, dialysed overnight against cohesin purification buffer 5 (50 mM $NaH_2PO4/Na_2HPO_4$ pH 7.5, 100 mM NaCl, 5 % glycerol, 0.5 mM TCEP), frozen in liquid nitrogen and stored at -80°C.

#### DNA synthesis

Supercoiled, closed-circle DNA mini-circles (MCs) were prepared using the ParA resolvase recombination system[5,6]. Upon induction of two plasmid-encoded enzymes, ParA and ITev, the ParA resolvase excises the target MC from the parental plasmid and the ITev endonuclease cleaves the vector backbone leading to its degradation. The parental vector, pRBPS_IVT7(2), was used to produce a 302 bp MC and additional plasmids (MC604, MC849, MC913, MC1217) were generated by inserting the required length of DNA between the ParA resolution sites to construct MCs of 604, 849, 913, and 1217 bp. An overnight culture, shaken at 28°C, of either pRBPS_IVT7 or its derivatives, in *E. coli* BL21 cells, was used to inoculate, at 1:100 dilution, 4 x 500 mL of LB (with 40 µg/mL kanamycin) and shaken at 37°C. When the OD600 reached 1.8, L-arabinose was added to 0.5% final concentration, to induce expression of ParA and ITev. After 4 hours of continued shaking at 37°C, cells were centrifuged and the pellets frozen. MC DNA was initially purified from these cell pellets using a NucleoBond Xtra Maxi kit (Machery-Nagel, 740414.50) per the manufacturer's protocol and the final pellet was resuspended in 10 mM TrisCl pH8, 1 mM EDTA (TE). To increase the final purity of the MC, the preparation was fractionated by differential precipitation using PEG6000. High MW DNA was initially precipitated by adding an equal volume of PEG6000 solution (14% PEG6000, 1.1 M NaCl, 1 mM EDTA). After incubation on ice (15 min), followed by centrifugation at 12,000 x g in a minifuge for 10 min, the supernatant was collected and adjusted to a final concentration of 15% PEG6000 then incubated on ice for 15 min. The precipitate (containing the MC) was pelleted and resuspended in TE. As a final step in the purification, T5 exonuclease (New England Biolabs) was added to digest any nicked or linearized DNA, leaving a preparation of closed-circle, supercoiled DNA MCs. During T5 exonuclease treatment, a restriction enzyme was added to allow selective digestion of any closed, circular DNA containing the recognition sequence of the restriction endonuclease. Accordingly, a restriction enzyme with no recognition sites in the MCs (BamHI-HF or HindIII-HF) was added during T5 exonuclease treatment to increase the efficiency of digestion of contaminating DNA. After the digestion, the enzymes were removed using the Monarch Clean-up kit (New England Biolabs) and eluted in TE, to generate a preparation of intact, closed-circle MCs. To linearize the MC, PstI digestion enzyme (Thermo Fisher) was added to the preparation, after which the enzyme was removed using Monarch Clean-up kit (New England Biolabs) and eluted in TE.

#### Mass photometry measurements

*Mass photometry data Acquisition and movie processing*

All mass photometry data were recorded using the TwoMP (Refeyn Ltd, Oxford). Measurements were done in silicon gaskets (GBL103250, Grace Bio-Labs) attached to precleaned microscope coverslips (24 × 50 mm, Menzel Gläser, VWR 630-2603), functionalised with (3-Aminopropyl)triethoxysilane (APTES, Sigma Aldrich, 440140). Coverslips were cleaned using three steps of 5 min bath sonication in water (Milli-Q® water, 18.2 MΩ·cm), 50% isopropanol in water and water again, and then dried under a stream of nitrogen gas. The coverslips were than treated with oxygen plasma for 3 min under 0.5 mbar of oxygen at 40% power (Zepto plasma cleaner, Diener Electronic). Immediately following plasma treatment, the coverslips were incubated for 3 min in a solution of 2% APTES in acetone (Sigma Aldrich, 34850-M). The coverslips were then washed in acetone and baked for 2 hours at 110 °C, followed by two cycles of bath sonication in 50% isopropanol in water and in Milli-Q water. The functionalised coverslips were then dried under a flow of nitrogen gas. Prior to the addition of the protein sample, buffer was added to the gasket and the focus position was adjusted to optimise the contrast at the glass-solution interface. For each measurement, a protein sample was loaded to the gasket containing the buffer and a 60 second movie was recorded using a field of view of 16.9 x 12.0 $mm^2$ and frame rate of 130 Hz (AcquireMP V2.5.0, Refeyn Ltd.). Movie processing and event detection was performed using DiscoverMP software (V2023 R1.2 Refeyn, Ltd.) where a rolling ratiometric movie was generated using a moving window of 5 frames, which corresponds to 38 ms integration time. For particles detection, threshold parameters (threshold 1 and 2) were set to their default values of 1.5 and 0.2, respectively. The measured contrasts were converted to mass using a contrast to mass protein calibrant as described previously[7,8].

##### *Histogram generation and partial concentrations calculations*

Mass histograms were generated a using custom-built python script. Mass histograms were generated with python package *scikit-learn*[9] using a kernel density estimator with a bandwidth of 5 kDa withing the detected mass range of 0 to 1500 kDa. The detected partial concentration of the molecular species was estimated by quantifying the relative peaks areas corresponding to each molecular mass. Quantification was done by fitting Gaussian functions to the measured mass histogram using python package *scipy.optimize*. The resulted areas were first normalised and then multiplied by the relevant measured total protein concentration as determined by absorbance to convert the relative detection of particles to their partial solution concentrations in a similar way as described previously[8,10,11]. These solution concentrations were then used to determine quantify the dissociation constant of an interaction, referred to as 'single-shot kd'. Histogram presentation was done using Origin software (OriginPro 2023, OriginLab)

##### *Histogram subtraction procedure and spectra reconstruction*

Since the Trimer sample is not perfectly homogenous and contains species with masses other than the intact Trimer (**Fig. S1,** see also[12]), the net Trimer concentration is lower than the total protein concentration measured. We also observed that the amount of the additional species found in the sample was not affected by the addition of STAG1, DNA or NIPBL, meaning that these unidentified biomolecular species do not participate in or affect the reactions we are quantifying.

To better visualize the peaks of the species of interest in the histograms we subtracted the constant background of molecular species that do not participate in the interactions of the cohesion holoenzyme from the experimentally observed distributions. Using the measurements of the individual components (STAG1, Trimer, NIPBL and DNA **Fig. S1** and **S2** for STAG1-Trimer example) we could isolate the contribution of the proteins of interest (the holoenzyme components) from the residual background of other biomolecules in the experimental histograms. To separate these two contributions, we first fitted a constraint Gaussian function to the peak of interest in the histogram of the individual components (for example the peak that correspond to the mass of the Trimer **Fig. S2** was constrained to 412 kDa $\pm$ 2% mass error). This Gaussian function then served as the line-shape for the contribution of this molecular specie to the full histogram. We then defined the rest of the histogram signal as a 'residual mass histogram', effectively splitting the signal obtained in the measurements of the individual components into two spectra, where each spectrum was normalized for its total area (alternatively, total number of counts) to be equal to 1. Examples for this step are shown in Fig. **S2**. This operation can be described by the following equation,

$$P_{component}^{Exp}(m) = G(\mu, \sigma; m) + R(m)$$

where, $P_{component}^{Exp}(m)$ is the experimental mass histogram of an individual component (STAG1, NIPBL, Trimer, DNA), $G(\mu, \sigma; m)$ is a Gaussian function correspond to the peak of interest and $R(m)$ is the residual.

Repeating this step for all the protein components, resulted in basic normalized probability densities corresponding to the peaks of interest, $P_i(m)$ where $i \in \{1,2,3,4\}$ ($P_i(m) = G(\mu_i, \sigma_i; m)$), representing the cohesin holoenzyme subunits (Trimer, STAG1, NIPBL) and the DNA and the corresponding residual spectra for each component ($R_i(m)$, where $i \in \{1,2,3,4\}$).

*Fitting the experimental histograms to quantify the partial concentrations.*

To extract the partial concentrations of the interacting species we used the following procedure. Each mass histogram that originates from a measurement containing more than 1 component (out of NIPBL, STAG1, Trimer, DNA) was normalised in a way that its total area (total number of counts) is equal to 1 ($P_{mixture}^{Exp}(m)$). Following the normalisation, the mass histogram was fitted to a modelled histogram constructed by a linear combination of the corresponding peaks of the individual components, their corresponding residuals and the peak(s) of their possible interactions, all are normalised to 1 and are weighted according to their mass fractions. The modelled histogram is therefore given by the following expression,

$$P_{mixture}^{Model}(m) = \sum_{i=1}^{n} f_i^p P_i(s_i m) + f_i^r R_i(s_i m) + \sum_{k=1}^{K} f_k^{inter} G_k(\mu_k, \sigma_k; m)$$

Here, $f_i^p$ and $f_i^r$ are the mass fraction of the *i*-th peak of interest, and the *i*-th residual, $s_i$ is a small scaling correction to represent the 2% variation in the contrast to mass calibration between different experiment and was set to the experimental limits of [0.98,1.02], $f_k^{inter}$ corresponds to the mass fraction of the *k*-th expected mass peak of the interacting subunits and $G_k(\mu_k, \sigma_k; m)$ represent the normalised Gaussian line-shape that correspond to the mass, $\mu_k$ and width, $\sigma_k$ of the detected complex. Knowledge of the best fitted mass fractions together with the initial protein concentrations in the mixtures enabled us to quantify the solution partial concentrations of the different subunits and their complexes from our mass histograms. To extract the vector of mass fractions, $\boldsymbol{F}$, we fitted our experimental results, $P_{mixture}^{Exp}(m)$, to the reconstructed histogram using non-negative least square algorithm (numpy.nnls) while adding the parameters $s_i, \mu_k, \sigma_k$ as additional parameters that were constrained to their expected experimental values with 2% variation resulting from the variations between experiments in the contrast to mass conversion and resolution.

*Presentation for the subtracted histograms*

For clarity of presentation, we show for the protein mixtures a subtracted histogram that emphasis the changes in the mass histogram that is directly related to the changes of the areas of the peaks of interest, which uncover the strengths of the examined interactions. (For example, Fig. 1c shows the subtracted histogram of STAG1 and Trimer interaction). To generate the subtracted presentation, we first perform the fitting procedure described above, and then subtracted from the experimental histogram the corresponding contributions of all the residual components, which corresponds to the following equation,

$$P_{mixture}^{Subtracted}(m) = P_{mixture}^{Exp}(m) - f_i^r R_i(s_i m)$$

For each experiment we show the resulted subtracted mass histograms of three technical repeats, their averaged histogram and the fitting residual that gives information on the goodness of fit. The residual was calculated by,

$$P_{mixture}^{Residual}(m) = P_{mixture}^{Exp}(m) - P_{mixture}^{Model}(m)$$

An example of a complete histogram data processing is shown in **Fig. S2.**

#### Experimental method for Holoenzyme interactions

To mimic biologically relevant conditions, we performed our experiments at concentrations comparable to those found inside the cell, which range from 70 nM for STAG1 to 300 nM for the Trimer's SCC1 subunit[13]. Unless stated otherwise, the proteins were incubated at 150 nM (total protein) for 5 minutes at room temperature and then 10-fold drop-diluted prior to data acquisition. The dilution was done directly in the silicon gasket that was prefilled with 22.5 mL of the same buffer solution, where the time interval between the dilution and the starting of data acquisition was approximately 10 seconds. The buffer use for incubation and measurement was 12.5 mM phosphate buffer pH 7.6, 25 mM NaCl, 1.25% glycerol and 2.5 mM $MgCl_2$ unless stated otherwise. When probing the effect of nucleotides, we added 2.5 mM ATP or 2.5 mM ATPγs to this buffer both during incubation and measurement. Prior to the measurements of mixtures, the different proteins were measured individually to verify serve as a baseline for further quantifications. For each mixture, at least 2 technical repeats were measured (Table S2).

*STAG1:Trimer high concentrations*

Experiments presented in **Fig. 1c** and **S2-3** were performed by incubating 150 nM STAG1 and 150 nM Trimer (Fig. 1c, top) and 80 nM STAG1 and 150 nM Trimer (Fig. 1c, bottom) Similar experiments were conducted (**Fig. S9a**) to examine the effect of ATP binding and hydrolysis on the STAG1-Trimer interaction.

*STAG1:Trimer titration at low nM concentration*

To validate that the measurements are performed at equilibrium, we confirmed that 5 minutes incubation is sufficient for complex assembly (on-rate) and that the complex does not disassemble over the course of a 1-minute video (off-rate, **Fig. S4**). Here, we mixed STAG1 and Trimer at 30 nM each, and followed the evolution of the mass histograms as a function of time. The mixture was measured at several time points following the mixing time, by transferring 10 mL of the sample into the measurement gasket that was prefilled with 10 mL of buffer. This dilution procedure resulted in the same final measurement concentration of 15 nM as when the proteins were incubated at higher concentrations, and validate that equilibrium is reached after 5 min of incubation and does not change over the course of a 60 sec data recording.

For better estimation of the reported dissociation constant, we performed a titration experiment at low nM concentrations (**Fig. 1d** and **S5-6**). Here, STAG1 concentration was varied between 3 and 30 nM, while keeping the Trimer concentration constant at 50 nM. For each titration concentration, three technical repeats were measured by taking 10 mL of the mixture and adding it to the prefilled measurement gasket resulting in an effective 2-fold dilution. We used 3 nM as lowest concentration of STAG1 in the titration, since that was the lowest concentration at which there was still a substantial and reproducible number of binding events when STAG1 was measured alone (599 ± 197 binding events, n = 3 technical repeats, quick decay over time due to protein sticking to the tube at nanomolar concentrations). The resulting mass histograms were fitted to a series of Gaussian functions to determine their solution partial concentrations (**Fig. S5b** and **S6**). Here we fitted the STAG1 and Trimer peaks, their interaction peak and the residual peaks to validate that both the total Trimer concentration as well as the residual peaks fraction stay constant throughout the titration (**Fig. S5b** and **Fig. S6**). To estimate the STAG1-Trimer dissociation constant we plotted the fraction of Trimer in complex with STAG1 as a function of free STAG1 (**Fig. 1D**, MP distributions shown in **Fig. S6**), and fitted to the Hill equation for 1 to 1 binding, $f_{STAG1-Trimer} = \frac{[STAG1]}{K_D+[STAG1]}$ using python package *scipy.optimize* by minimizing the square difference between the data and the model.

*Trimer: NIPBL and NIPBL:STAG1 interactions*

The experiments for the quantification of the interaction between NIPBL/DN-NIPBL and the Trimer (**Fig. S9, Fig. S10, S16**) as well as the possible interaction between STAG1 and NIPBL (**Fig. S9, Fig. S10**) were performed using identical procedure for the STAG1-Trimer experiments described above, where 150 nM of NIPBL was mixed with 150 nM of the Trimer or STAG1.

*Holoenzyme interactions*

To follow the formation of the full holoenzyme, we followed identical experimental procedure and mixed 150 nM of each protein (STAG1, NIPBL and Trimer) with and without the addition of ATP or ATPgS (**Fig. 1e** and **Fig. S9, Fig. S10**).

#### Proteins - DNA interaction

Experimental results involving the interaction of the protein subunits with the minicircular DNA were perform by mixing equal concentrations of protein and DNA at 150 nM (**Fig. 2** and **3a,b**). The nucleotides were first added to the solution containing DNA, which was later mixed with the protein solution. A similar procedure was followed for the interactions of the protein complexes (STAG1-Trimer and full holoenzyme) with DNA (**Fig. 3d**, **Fig. S8, Fig. S9, Fig. S10**). Here, we first mixed the protein subunits with each other and following 3 minutes of incubation we measured the resultant mass histogram to validate that the protein-protein interactions were equilibrated. We then added DNA to reach equimolar concentrations of 150 nM for all the components and allowed 3 minutes additional incubation

*Testing the effect of the buffer and the effect of ionic strength on the STAG1-Trimer-DNA interaction*

To exclude the possibility that the phosphate buffer affects the affinity of the STAG1-Trimer complex for DNA in the presence of ATP or ATPgS, we repeated these experiments in different buffers (**Fig. S13** and **Fig. S14**, top two panels in both). Here, we repeated the same experimental procedure detailed above, but replaced the incubation and measurement buffer from 12.5 mM phosphate buffer to 20 mM Tris-HCl buffer at pH 7.6. In this buffer the protein concentrations were identical to those observed in our standard phosphate buffer at equimolar concentration of 150 nM. In addition, the resulting mass histograms were similar to those obtained in the phosphate buffer.

To test the effect of the ionic strength on the STAG1-Trimer interaction with DNA, we mixed 150nM STAG1 with 150nM of Trimer in a buffered solution containing 20 mM HEPES pH 7.6, 50 mM NaCl, 2 mM $MgCl_2$ and 1.25% glycerol and then added 150 nM MC302. The additional contribution to the ionic strength of 25 mM NaCl is comparable to that of 2.5 mM of nucleotide and therefore serves as a control for the effect of ionic strength on STAG1-Trimer-DNA interactions in the presence of nucleotides. These results show that an increase of the ionic strength of the buffer weakens the interaction between the STAG1-Trimer complex and DNA (**Fig. S13** and **Fig. S14**, bottom panel in both).

### Supplementary Figures

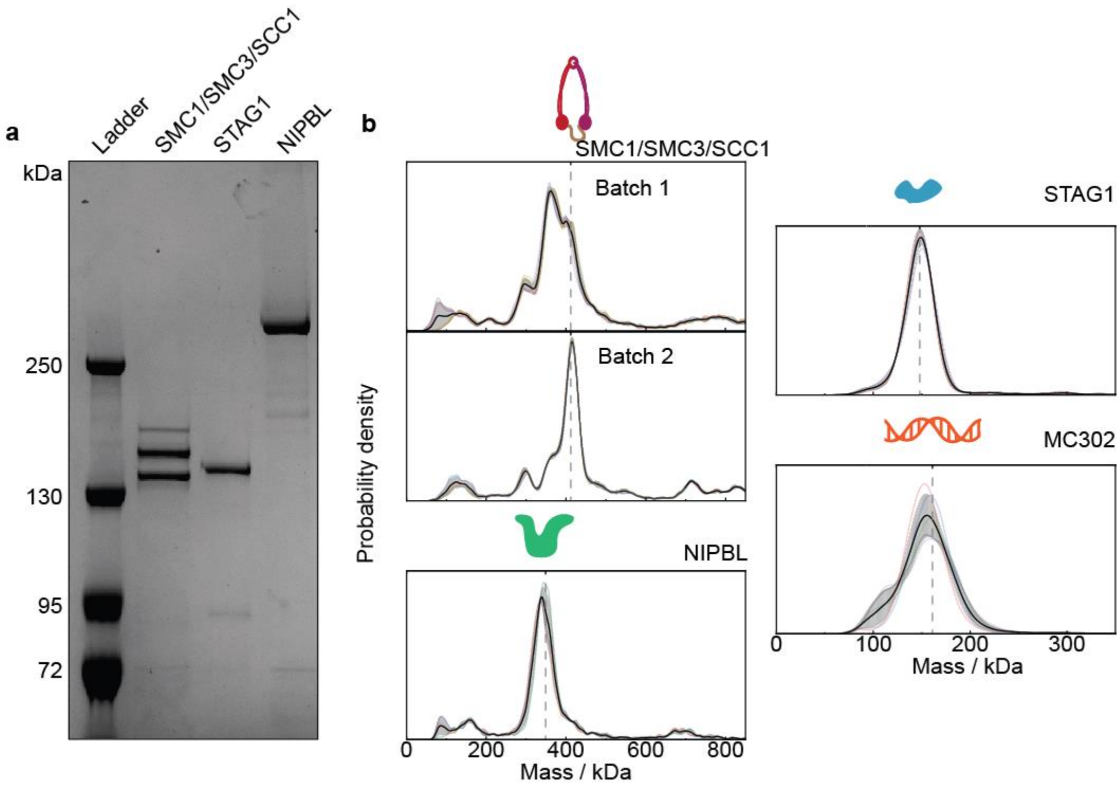


**Figure S1. MP distributions for the SMC1/SMC3/kleisin Trimer, NIPBL, STAG1, and MC302 DNA. a** Coomassie blue SDS-PAGE denaturing gel (6% Tris-Glycine) confirms successful purification of the SMC1/SMC3/SCC1 Trimer, STAG1, and NIPBL and reveals minor impurities. **b** Each panel shows the average (black) ± standard deviation (grey) of 3 or more technical repeats (solid colored lines), and the position of the monomeric protein of interest (vertical dashed grey). The fractions of the number of counts that constitutes to our biomolecule of interest relative to the total number of detected biomolecules in that sample correspond to: 22% (SMC1/SMC3/SCC1 Trimer batch 1), 41% (SMC1/SMC3/SCC1 Trimer Batch 2), 49% (NIPBL), 92% (STAG1), and 76% (MC302). The average number of detected particles in the 0-1500 kDa range for the technical repeats are: $7.2\times10^3$ (SMC1/SMC3/SCC1 Trimer batch 1, n = 3), $9.7\times10^3$ (SMC1/SMC3/SCC1 Trimer batch 2, n = 5), $3.5\times10^3$ (NIPBL, n = 3), $5.4\times10^3$ (STAG1, n = 4), $7.9\times10^3$ (MC302, n = 5)

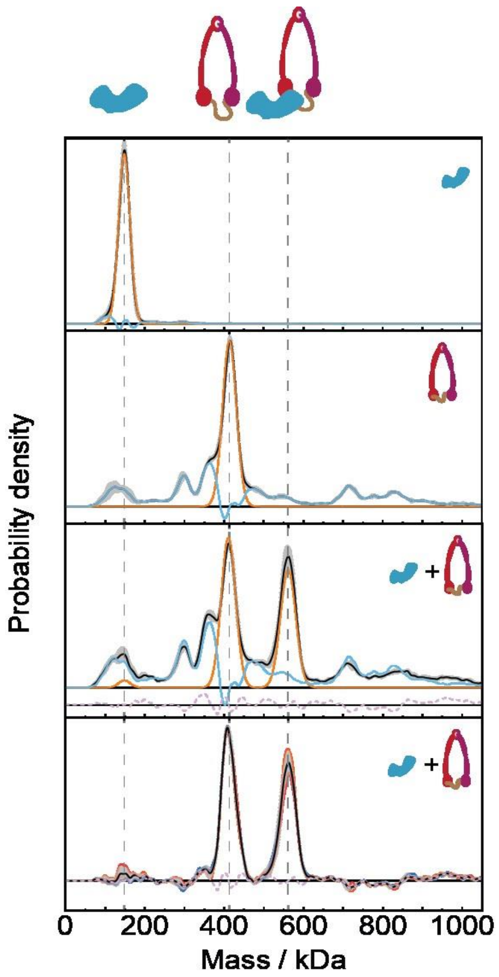


**Figure S2. Procedure for subtraction residual signal that does not participate in the reaction**

The procedure for residual subtraction starts with identification of the peak of interest (orange) and residual (blue) in the individual components when they are measured alone (top two panels). Then the mean experimental distribution from the mixture (black line) is reconstructed using the residual of the individual components (blue), their peak of interest, and any possible interaction peak(s) (both orange, third panel). We allow for a horizontal scaling of 5% of the entire experimentally measured spectrum and a scaling of 2% for the component and interaction peaks to account for differences in contrast to mass conversion between different measurements. The difference between the experimental measurement and the reconstruction is shown on a separate axis for clarity (dotted pink line). Having identified the contribution of the residuals in the average experimental distribution, we can subtract these from each technical repeat to generate the experimental distribution with residuals removed (bottom panel). Plots shown is from the same experiments as shown in Fig. 1C, Fig. S1, and Fig. S3.

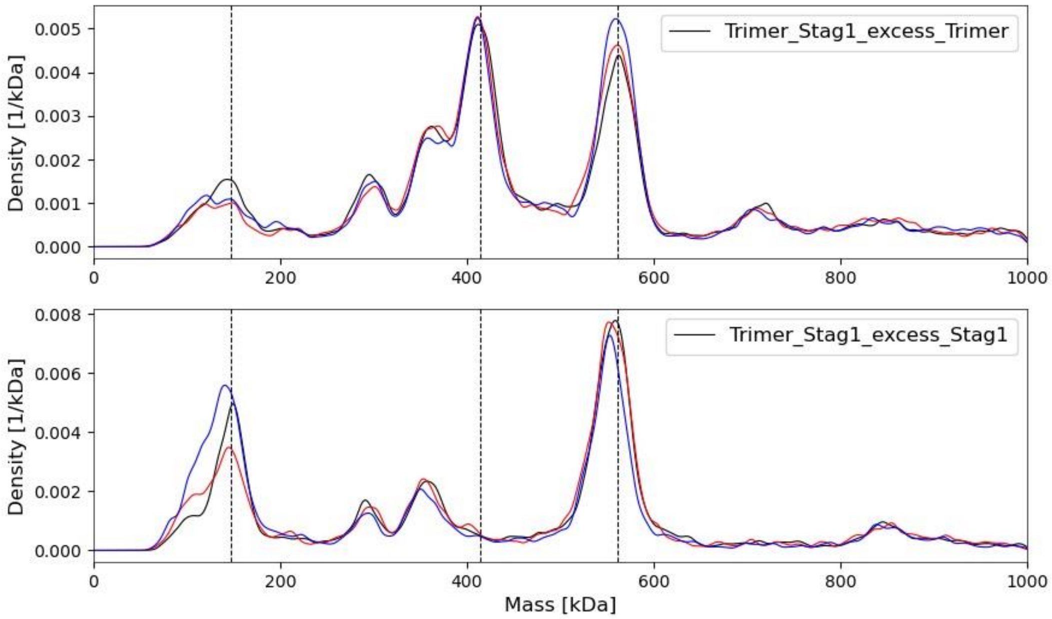


**Figure S3. Mass distributions for the STAG1-Trimer plots shown in Fig. 1C without subtraction.**

The figure shows the unprocessed normalised experimental mass histograms of a mixture of 80 nM STAG1 and 150 nM Trimer following 5 min of incubation at room temperature. Each panel show 3 technical repeats within the mass range of 0-1000 kDa. Vertical lines correspond to the expected masses of STAG1, Trimer and their complex. The average number of detected particles for the experiments in the 0-1500 kDa range for the technical repeats are $4.7\times10^3$ (Excess Trimer, n = 3), and $9.8\times10^3$ (Excess Trimer, n = 5).

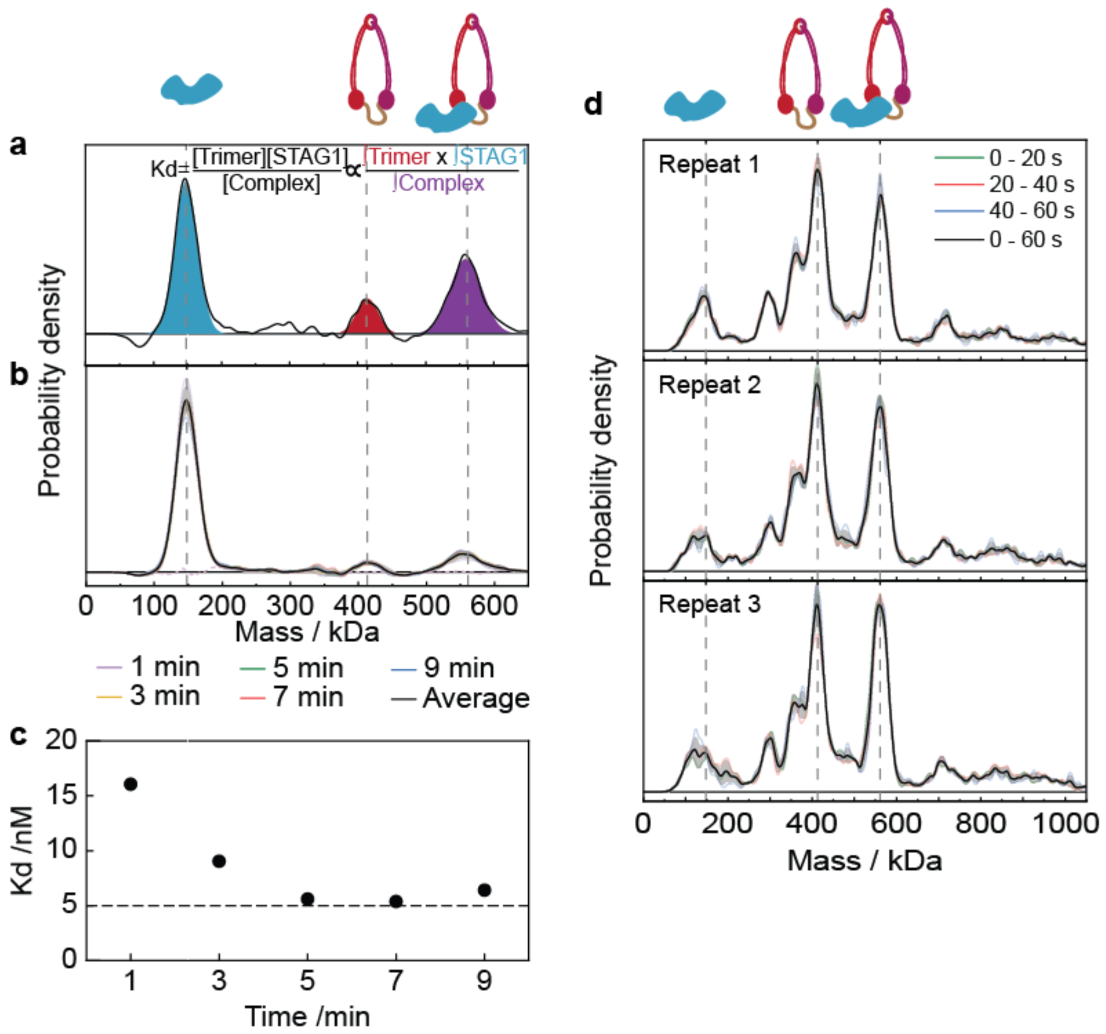


**Figure S4. Validation of the on and off-rate of the STAG1-Trimer interactions.**

**a**, To calculate the dissociation constant from a single measurement (single-shot $K_D$), the peaks of interest are fitted with a Gaussian, with blue, red and purple filled peaks corresponding to the fits for the STAG1, Trimer and STAG1-Trimer complex mass peaks. The area of a fitted Gaussian function corresponds to the solution concentration of the corresponding species (**Fig. S2A**, Soltermann et al. 2020, Fineberg et al. 2020). These numbers are then scaled by a factor to consider the incubation concentration and a correction factor to convert MP counts to molarity. **b**, STAG1 + Trimer measured 1, 3, 5, 7, and 9 minutes after mixing to probe the on-rate of the interaction. Proteins were mixed at 30 nM each and then two-fold diluted into the gasket directly prior to taking the measurement. Each measurement contains at least $4.1\times10^3$ detected particles in the 0-1500 kDa range. **c,** Single-shot dissociation constant of these measurements show a plateau after 5 minutes with a single-shot $K_D$ of ~ 5 nM, providing an estimate of the on-rate at these concentrations (minutes) and showing that an incubation of 5 minutes is sufficient for complex assembly (especially considering that for all other experiments except the titration, the biomolecules were mixed at 150 nM each and then diluted 10-fold prior to a measurement). **d,** Three technical repeats of STAG1-Trimer (same measurement as shown in **Fig. S2** and **Fig. 1C**), with each measurement split into three temporal windows. Since the distributions per repeat show neat agreement (the STAG1-Trimer complex abundance does not change over the course of a measurement), the off-rate at these concentrations (15 nM per protein) is sufficiently slow (> 1 minute) to not significantly affect the measurement.

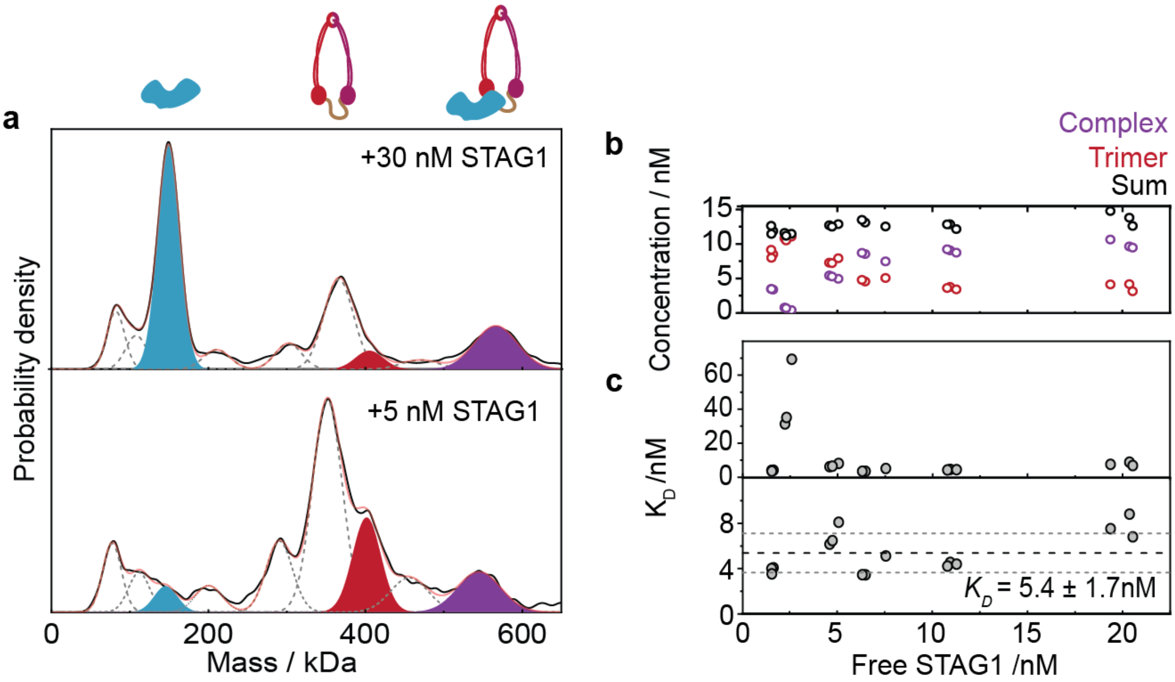


**Figure S5. Procedure for dissociation constant quantification from individual measurements**

**a,** Representative mass histograms for two measurements in the titration. The peaks of STAG1 (blue), the Trimer (red), and STAG1-Trimer (purple) are indicated. **b**, Quantification of the concentration of Trimer-STAG1 complex (purple), free Trimer (red), and total Trimer (black) for the STAG1 titration (Fig. 1C, Fig. S6). As expected, the concentration decrease of free Trimer with increasing amounts of STAG1 is on par with the increase the STAG1-Trimer concentration increase, and the total Trimer content stays constant (12.6 ± 0.9 nM). **c**, Single-shot dissociation constants for the Trimer + STAG1 titration showing all datapoints (top panel), or with the three outliers (all technical repeats of a single measurement) removed. The mean of the measurements in the bottom panel (black dashed line) and the standard deviation (grey dashed line) are indicated. The single-shot $K_D$ determined in this manner agrees neatly with the one from the fitted hill curve (Fig. 1C). Calculation of the average $K_D$ value for all the datapoints without excluding the outlier gives 12 ± 16 nM.

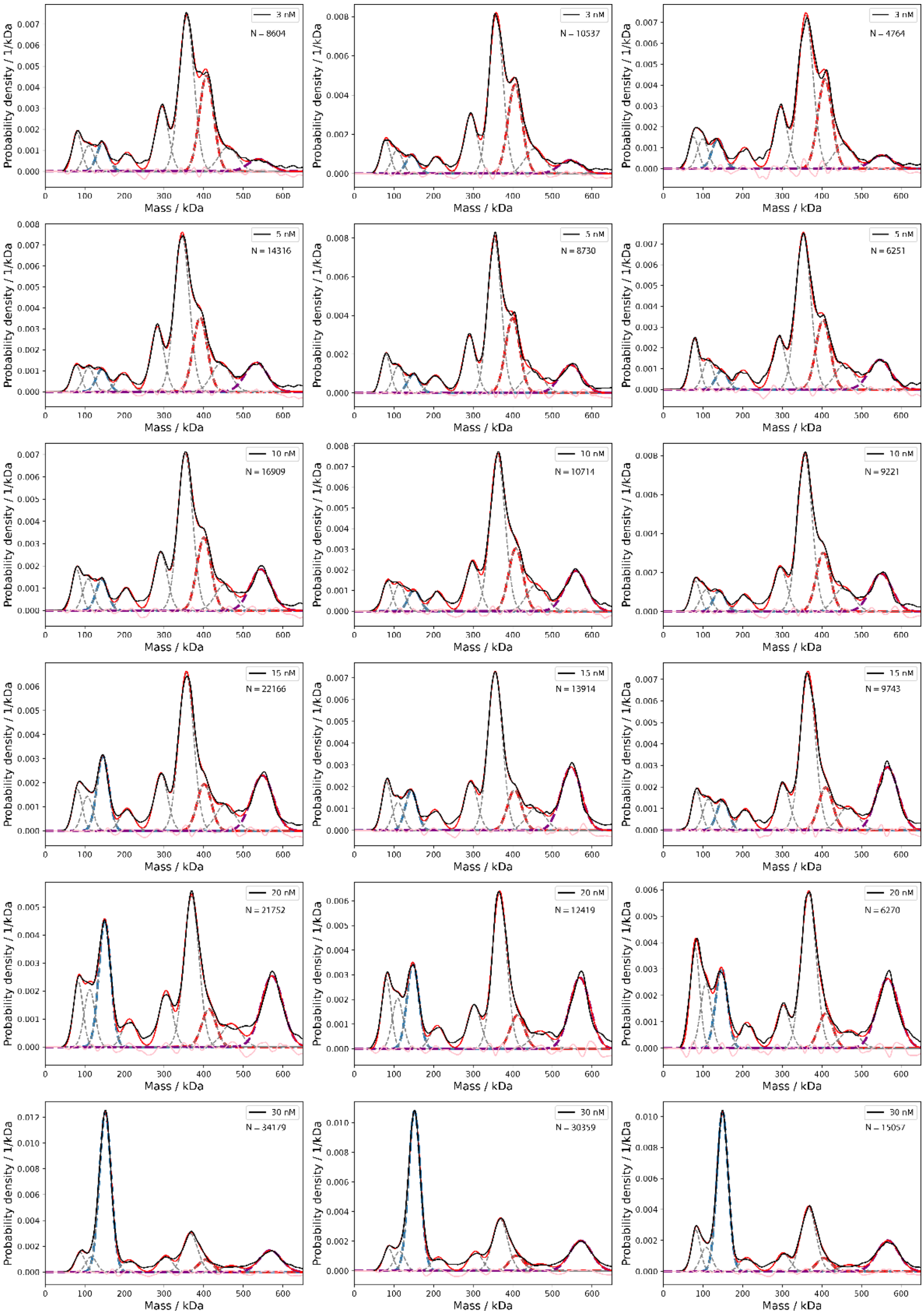


**Figure S6. Mass distributions for the STAG1-Trimer titration.**

Individual mass distributions for the datapoints shown in **Fig. 1d**. The experimental distribution (solid black), the Gaussian fits to the STAG1 peak (dashed blue), Trimer (dashed orange), STAG1-Trimer (dashed purple), and the residual signal (dashed grey) are shown. The green line shows the difference between the sum of all Gaussian fits (red) and the experimental distribution. The Trimer concentration was fixed at 50 nM for all measurements while the concentration of STAG1 was varied between 3 and 30 nM as indicated for each histogram. Three technical repeats were measured at each concentration. The measured number of particles for the construction of each histogram is shown in each panel.

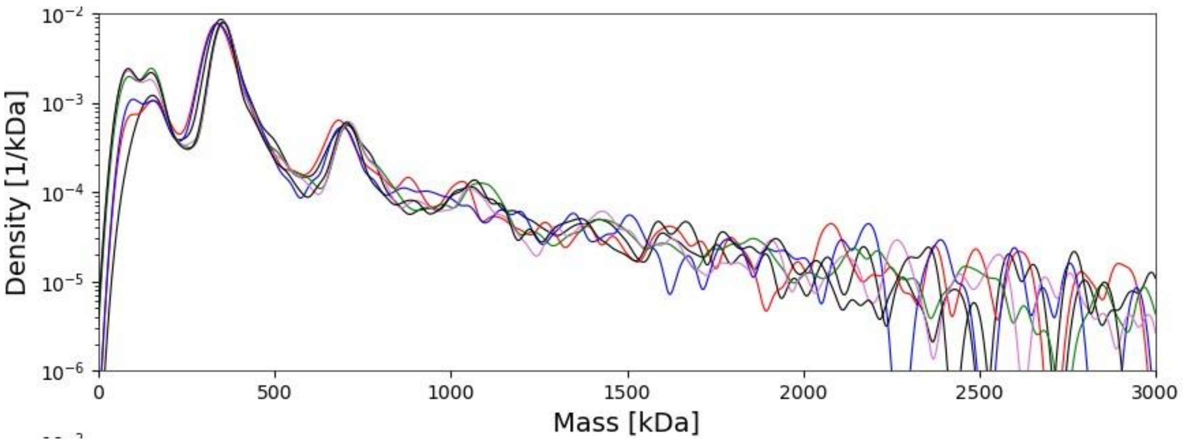


**Figure S7. NIPBL mass on logarithmic scale reveals self-oligomerization.**

Measured mass distribution of NIPBL incubated at concentration of 150 nM and room temperature for approximately 5 min. The sample was measured using the fast dilution method explained in the Method section. The results show that the most abundant state of the protein is the monomer, however self-oligomerisation takes place under the measured conditions. Each technical repeat has at least $2.5\times10^3$ detected particles in the 0-3000 range (n=6).

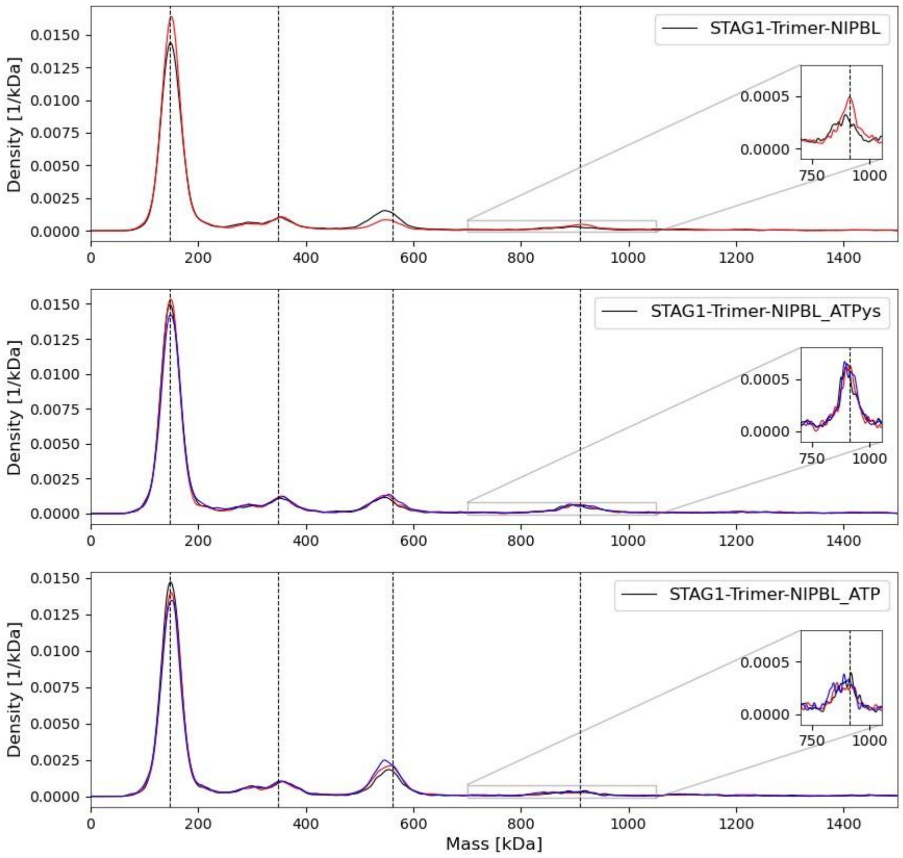


**Figure S8. Mass distributions for the STAG1-Trimer-NIPBL plots shown in Fig. 1E without subtraction.**

Unprocessed normalised experimental mass histograms of a mixture of 150 nM STAG1, 150 nM Trimer and 150 nM of NIPBL. All concentrations are given for the total protein content in each sample. Measurements were conducted following 5 min of incubation at room temperature in similar buffered solution with or without the addition of 2.5 mM of ATP and ATPgS, as indicated for each panel. Each panel show 3 technical repeats within the mass range of 0-1500 kDa. Insets show the mass range corresponding to the mass of the holoenzyme on expanded scale. Vertical lines correspond to the expected masses of STAG1, Trimer, STAG1-Trimer complex and the fully assembled holoenzyme. The average number of detected particles for the experiments in the 0-1500 kDa range for the technical repeats are $2.7\times10^4$ (APO, n = 2), $1.1\times10^4$ (ATPγs, n = 3), and $1.0\times10^4$ (ATP, n = 3).

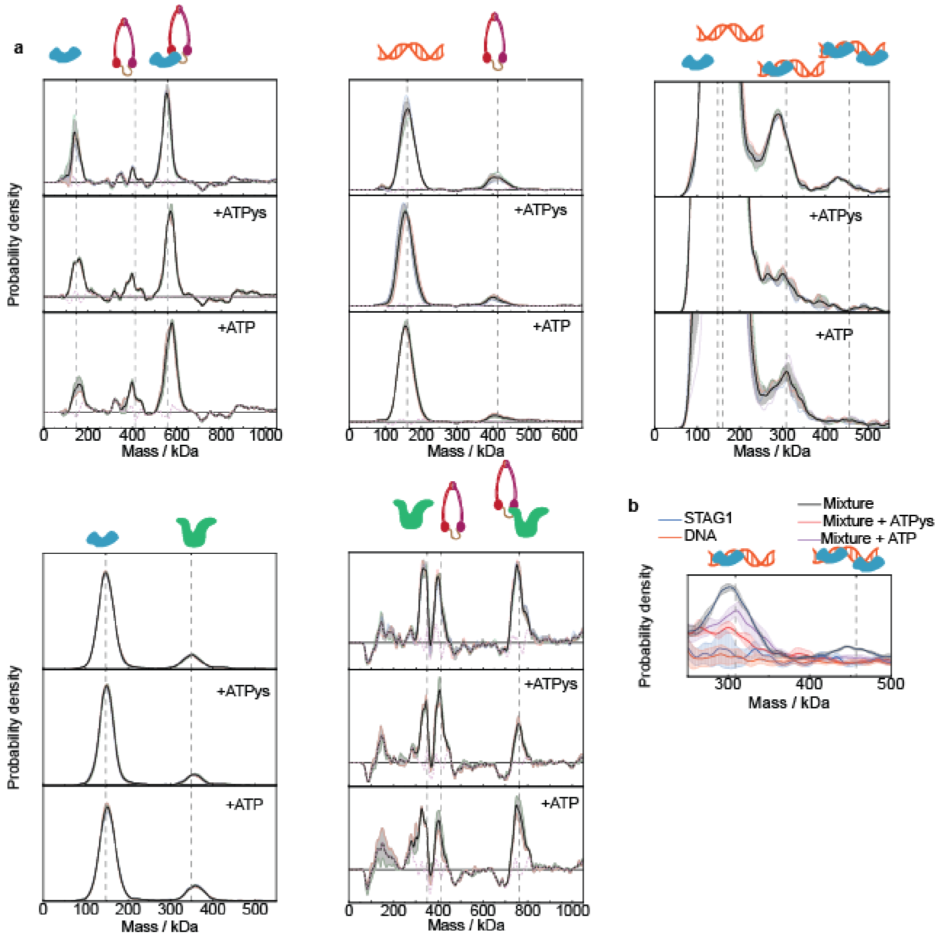


**Figure S9. The one-to-one interactions show no dependence on ATP binding or hydrolysis.**

**a**, MP distributions for various mixtures in APO state (top), +ATPγs (middle), or +ATP (bottom). The expected positions of the peaks (vertical grey dashed lines), technical repeats (colored lines), mean (black), and standard deviation (grey) are depicted. The residuals were subtracted from measurements including the Trimer (because it has a large residual signal) and for these the error of the reconstruction is shown (pink dashed line). NIPBL-Trimer and STAG1-Trimer interact strongly irrespective of nucleotide presence, whereas Trimer-DNA and STAG1-NIPBL do not interact at all. The STAG1-DNA is already very weak in the APO state, which makes reproducible detection challenging. The average number of detected particles for the experiments in the 0-1500 kDa range for the technical repeats are $9.2\times10^3$ (STAG1-Trimer APO, n = 3), $4.6\times10^3$ (STAG1-Trimer ATPγs, n = 2), $6.7\times10^3$ (STAG1-Trimer ATP, n = 2), $5.8\times10^3$ (Trimer-DNA APO, n = 3), $1.7\times10^4$ (Trimer-DNA ATPγs, n = 3), $1.3\times10^4$ (Trimer-DNA ATP, n = 3), $2.0\times10^4$ (STAG1-DNA APO, n = 3), $6.9\times10^3$ (STAG1-DNA ATPγs, n = 3), $8.7\times10^3$ (STAG1-DNA ATP, n = 4), $9.1\times10^3$ (STAG1-NIPBL APO, n = 3), $3.5\times10^3$ (STAG1-NIPBL ATPγs, n = 3), $5.7\times10^3$ (STAG1-NIPBL ATP, n = 3), $4.2\times10^3$ (NIPBL-Trimer APO, n = 3), $5.8\times10^3$ (NIPBL-Trimer ATPγs, n = 2), and $5.6\times10^3$ (NIPBL-Trimer ATP, n = 2). **b**, Zoomed distribution for the STAG1 + DNA mixture in the region of the interaction peaks for the 3 measurements shown in (a) (black, red, and purple) and STAG1 (blue) and DNA (orange) when measured alone. For the mixtures we see an enrichment in the peak region for nucleotide conditions, which we interpret the STAG1-DNA interaction not being significantly affected by nucleotide presence (as expected, because STAG1 does not have an ATPase). Full distributions are reported in Fig. S1 and this figure.

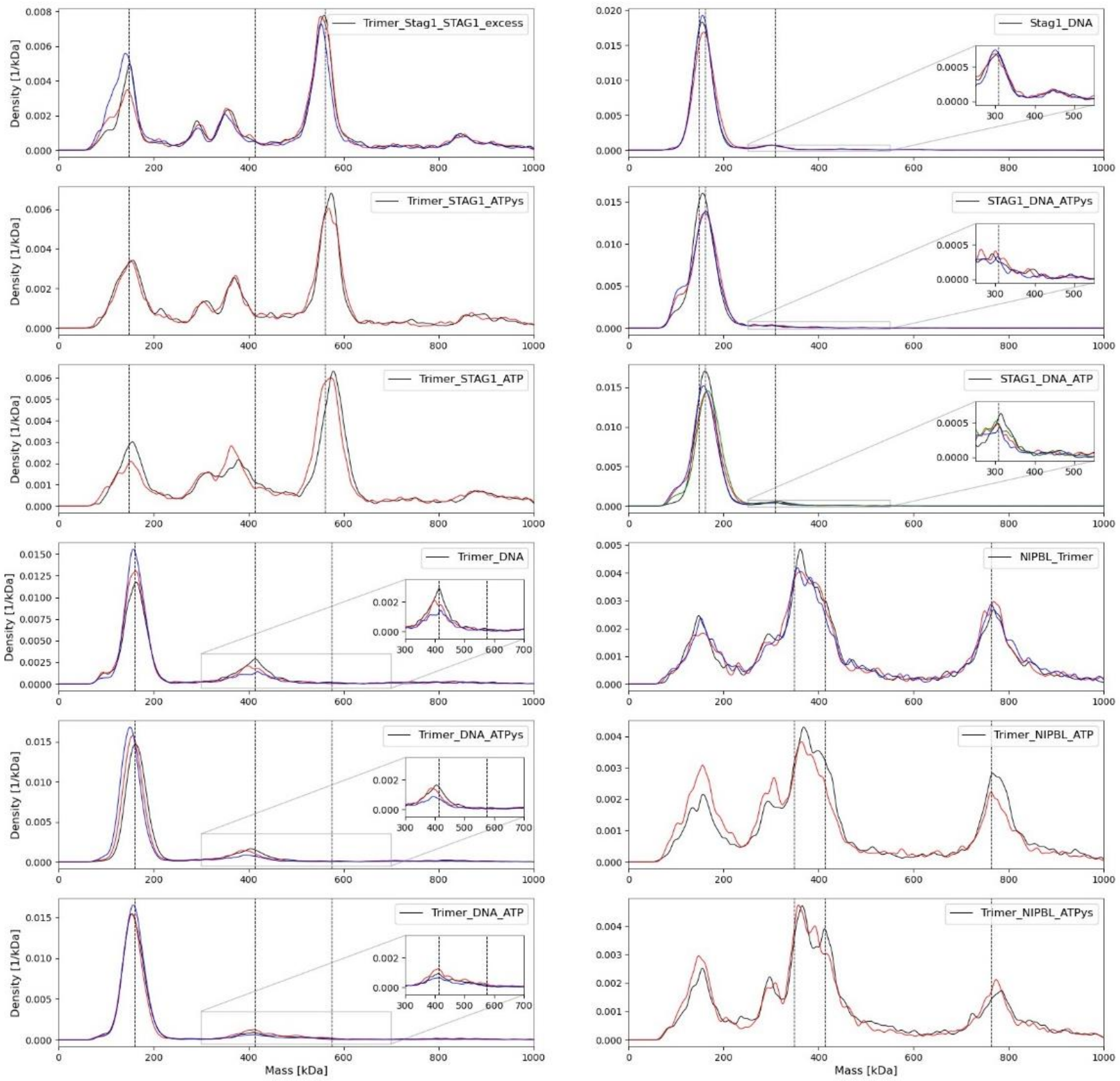


**Figure S10. Raw mass distributions for all measured 1:1 interactions.**

This figure shows the unprocessed normalised experimental mass histograms of a mixture all the examined 1:1 interactions in this work between STAG1, Trimer, NIPBL and DNA (subtracted distributions shown in Fig. S9, where applicable. The mixed biomolecules are indicated at the top right corner for each panel together with the type of nucleotide, if added (see Methods for buffers conditions). All proteins were mixed at equimolar concentration of 150nM. All the mass histograms are shown before the residual subtraction procedure was applied, for each measurement at least 2 2 technical repeats were measured, indicated by different colors. Vertical lines indicate the expected masses of the mixed subunits and the expected mass of their complex (schematics depicted in Fig. S9). Insets, show the mass range were the interaction peak is expected.

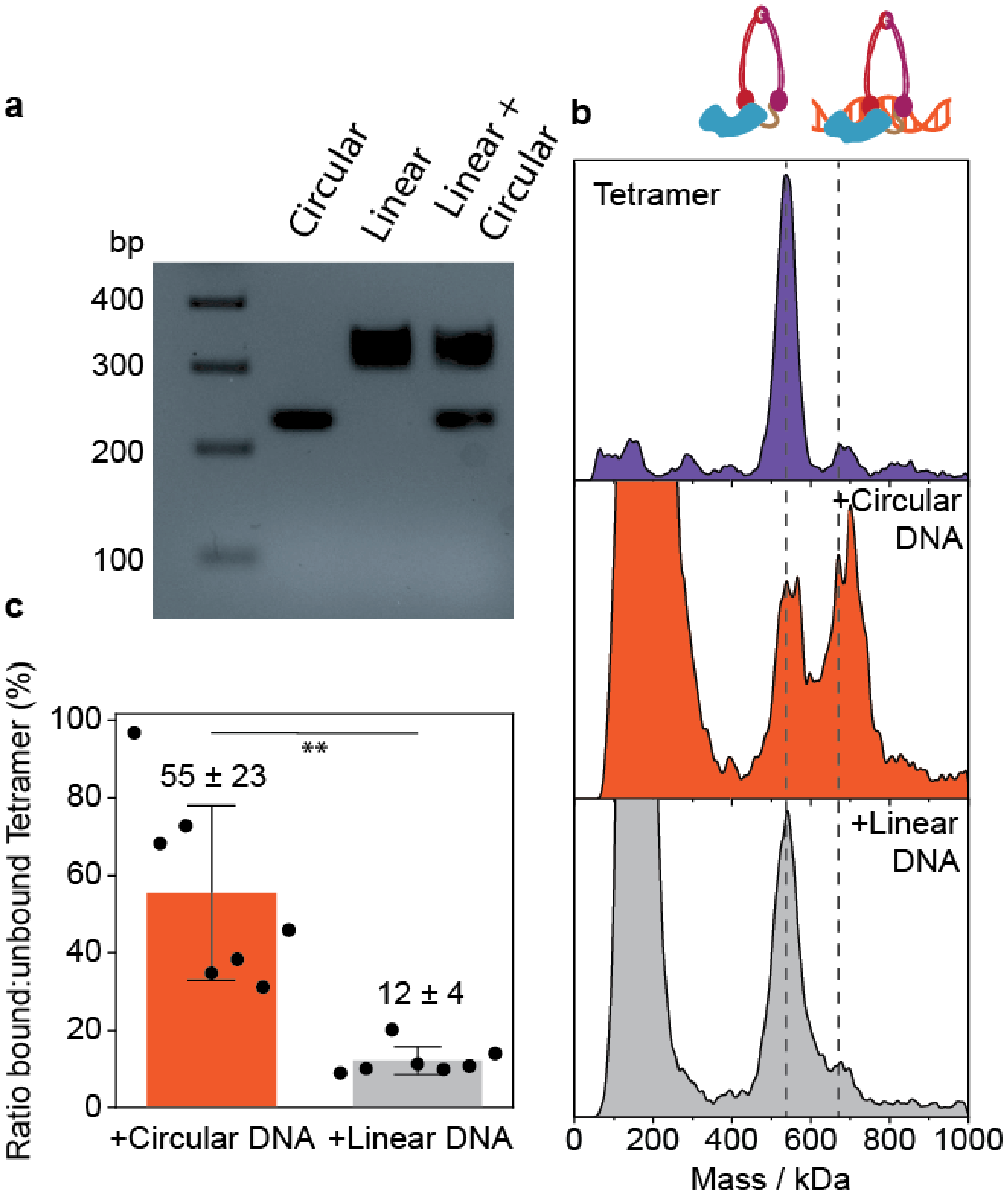


**Figure S11. STAG1-Trimer binding to circular and linear DNA.**

**a** 2% agarose gel showing circular MC302 and linear MC302 by digestion with PstI. Ladder is double stranded linear DNA and staining with Ethidium Bromide. Although equal amounts of linear and supercoiled DNA, as determined by absorbance (260 nm), were loaded in the last lane, the digested sample intercalates more of the fluorescent dye, so it appears more abundant. The faster migration of the circular DNA than the linear DNA suggests the circular DNA is supercoiled[14] **b** Representaive unprocessed normalised experimental mass histograms of the STAG1-Trimer (Tetramer, top, purple), the Tetramer incubated with Circular MC302 DNA (middle, orange), and the Tetramer incubated with linearized MC302 DNA (grey, bottom). Vertical lines indicate the expected masses of the mixed subunits and the expected mass of their complex. The average number of detected particles for the experiments in the 0-1500 kDa range for the technical repeats are 5600 (Tetramer, n = 3), 19000 (Tetramer + Circular MC302, n = 7), 30000 (Tetramer + Linear MC302, n = 7). **c** Quantification of the ratio of the Tetramer-DNA peak to the free Tetramer peak. As the free Tetramer also has counts in the expected mass region of the Tetramer-DNA peak, this background signal was subtracted from the histograms of the mixtures. Errorbars indicate the standard deviation from seven technical repeats, which were pooled from two different days. **: p< 0.005

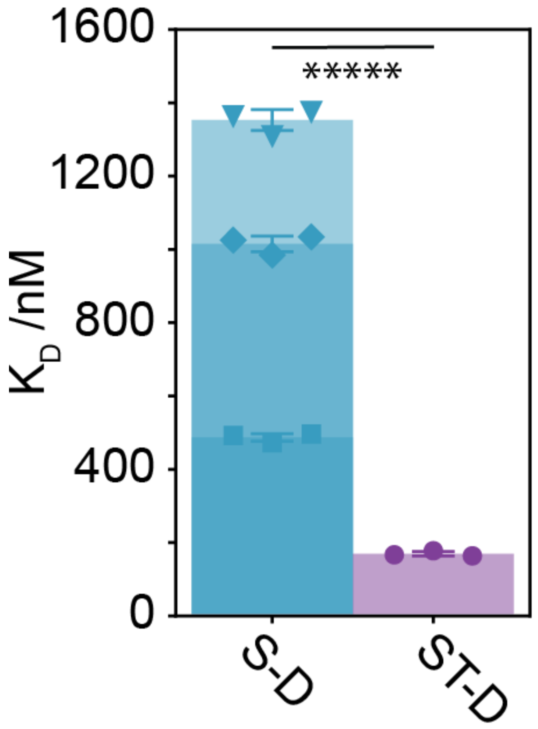


**Figure S12. Interaction of STAG1 and STAG-Trimer with DNA.**

Since the STAG1 & DNA peaks overlap (Fig. 2C, left peak), we cannot fit them separately and instead assume various ratios of STAG1 : DNA in that peak: 50:50 (triangles, top, shown in main, $K_D$ 1353 ± 29 nM), 75:25 (or 25:75) (diamonds, middle, $K_D$ 1015 ± 22 nM), or 90:10 (or 10:90) (squares, bottom, $K_D$ 487 ± 10 nM). Since the proteins were mixed in a ratio that gives equal number of counts when measured separately, we deem the 50:50 approximation reasonable. *****:p<0.000005 for all cases.

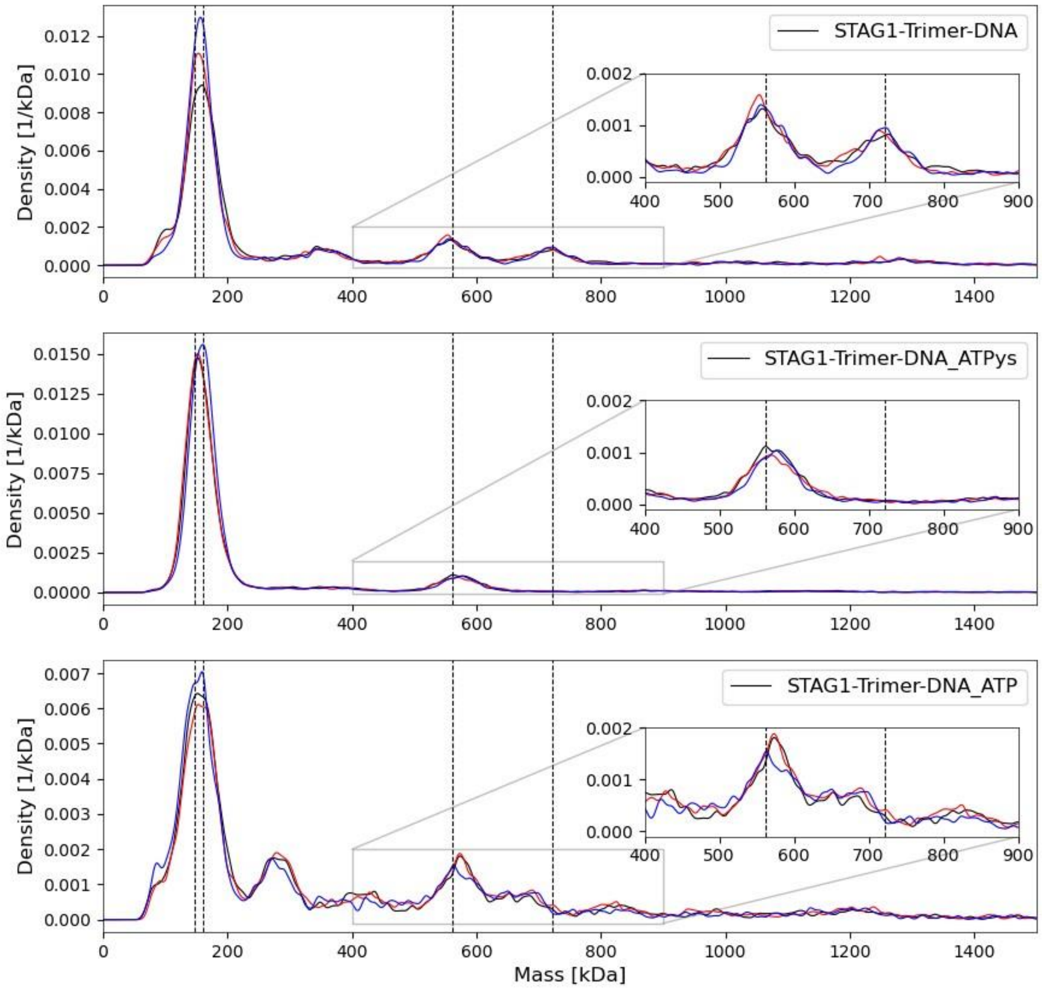


**Figure S13. Normalised mass distributions for the STAG1-Trimer-DNA plots shown in Fig. 3A.**

Unprocessed normalised experimental mass histograms of a mixture of 150 nM STAG1, 150 nM Trimer and 150 nM of DNA (total protein concentrations) with and without the addition of 2.5 mM of nucleotides. Each panel show 3 technical repeats within the mass range between 0 and 1500 kDa. Insets show the mass range corresponding to masses of the expected STAG1-Trimer complex and its interaction with DNA on expanded scale. Vertical lines indicate the expected masses of the mixed subunits and the expected mass of their complex (schematics depicted in Fig. 3a). The average number of detected particles for the experiments in the 0-1500 kDa range for the technical repeats are $6.7 \times 10^3$(STAG1-Trimer-DNA APO, n = 3), $1.6 \times 10^4$ (STAG1-Trimer-DNA ATPγs, n = 3), and $4.6 \times 10^3$ (STAG1-Trimer-DNA ATP, n = 3).

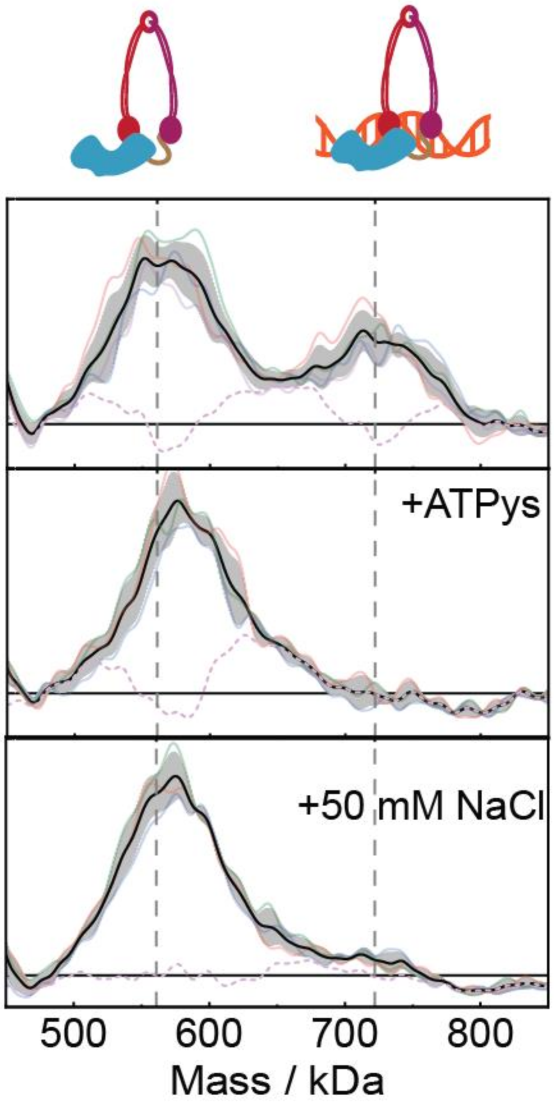


**Figure S14. STAG1 + Trimer + DNA measurements in a different buffer.**

The top two panels are measured in 20 mM Tris pH 7.6, 25 mM NaCl, 2 mM $MgCl_2$, 1.25% glycerol (+ ATPγs) and show the same trend as the measurements in the standard buffer (12.5 mM $NaPO_4$ pH 7.6, 25 mM NaCl, 1.25% glycerol, 2 mM $MgCl_2$, Fig. 3A), thus ruling out the possibility that the Trimer-STAG1 release is influenced by phosphate presence in the standard buffer. The bottom panel was measured in 20 mM HEPES, 50 mM NaCl 2 mM $MgCl_2$, 1.25% glycerol, hence the absence of a Trimer-STAG1-DNA peak shows that electrostatic contributions are important for this interaction. The average number of detected particles for the experiments in the 0-1500 kDa range for the technical repeats are $1.5 \times 10^4$ (APO APO, n = 4), $1.7 \times 10^4$ (ATPγs, n = 2), $2.1 \times 10^4$ (+50 mM NaCl, n = 3).

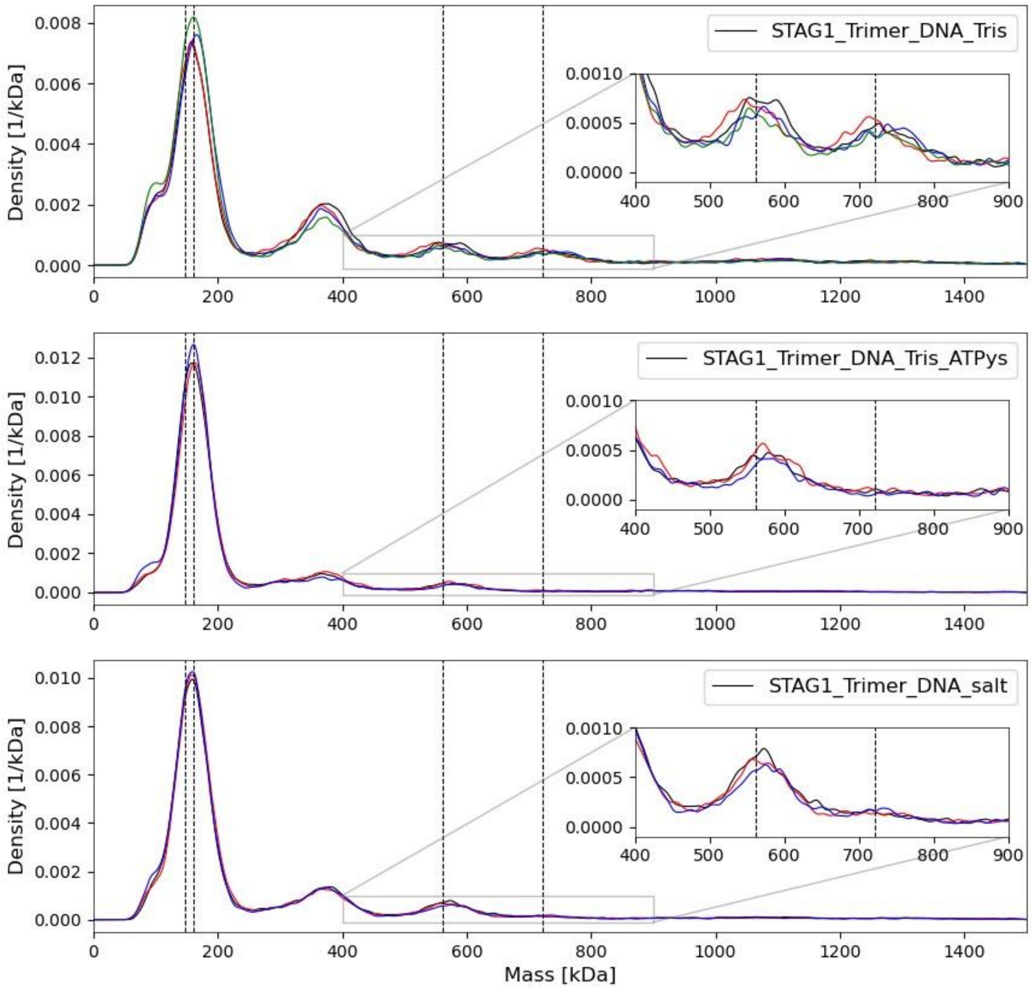


**Figure S15. Mass distributions for the STAG1-Trimer-DNA plots shown in Fig. S13 without subtraction.**

This figure shows the unprocessed mass histogram of mixing STAG1, Trimer and DNA at equimolar concentration of 150 nM at the same buffers conditions as indicated in **Fig. S14**. Vertical lines correspond to the expected masses of DNA, STAG1, Trimer, Trimer-STAG1 complex and the complete complex STAG1-Trimer-DNA. Insets show the mass range that corresponds to the messes of the STG1-Trimer and STAG1-Trimer-DNA complexes. Between 3 and 4 technical replicates were measured at each solution condition, indicated by the different colors.

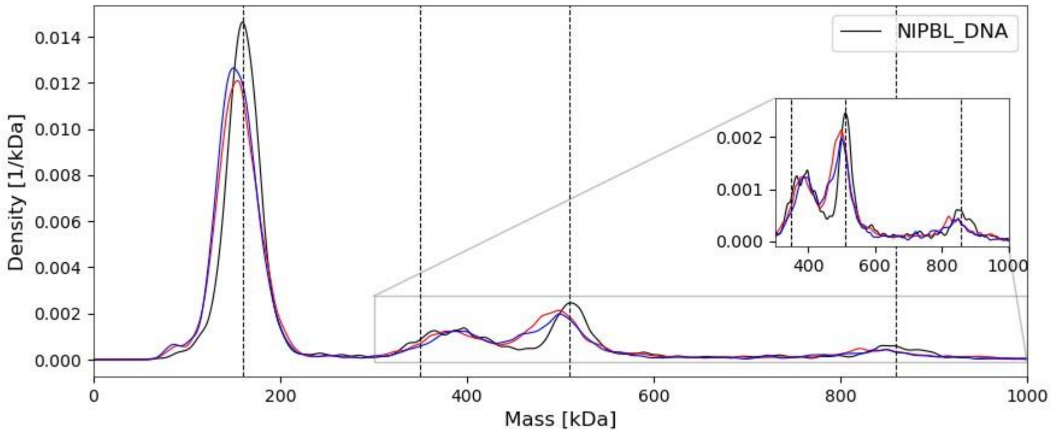


**Figure S16. Mass distributions for the NIPBL-DNA plots shown in Fig. 3C without subtraction.**

The figure shows the unprocessed normalised experimental mass histograms of a mixture of 150 nM NIPBL and 150 nM DNA following 5 min of incubation at room temperature. The 3 technical repeats are shown with different colors. Vertical lines correspond to the expected masses of DNA, NIPBL and their 1:1 and 2:1 NIPBL:DNA complexes. The average number of detected particles for the experiments in the 0-1500 kDa range for the technical repeats is $7.7 \times 10^3$(n=3).

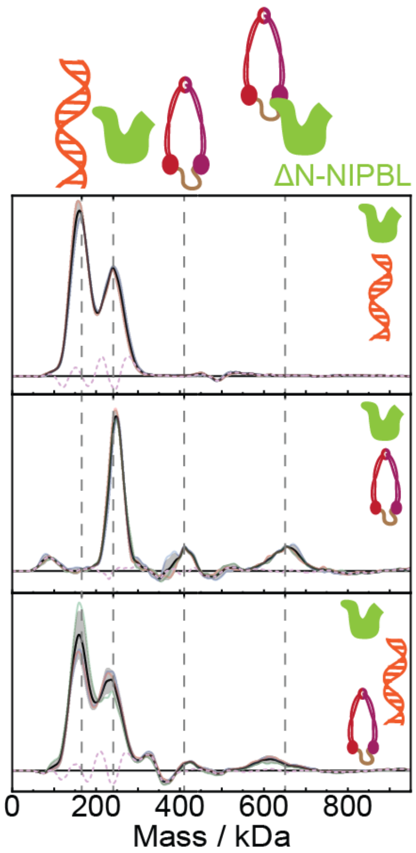


**Figure S16. N-terminal Truncated-NIPBL binds the Trimer, but not DNA.** Top: Truncated NIPBL mixed with DNA, does not show an interaction peak. Middle: Truncated-NIPBL does bind the Trimer. Bottom: Truncated NIPBL-Trimer complex does not bind DNA. Each panel shows the average (black) ± standard deviation (grey) of 3 or more technical repeats (solid colored lines), the position of the monomeric protein of interest (vertical dashed grey). The average number of detected particles for the experiments in the 0-1500 kDa range for the technical repeats are $2.2 \times 10^4$ (ΔN-NIPBL-DNA, n = 3), $1.3 \times 10^4$ (ΔN-NIPBL-Trimer, n = 3), and $2.2 \times 10^4$ (ΔN-NIPBL-DNA, n = 3).

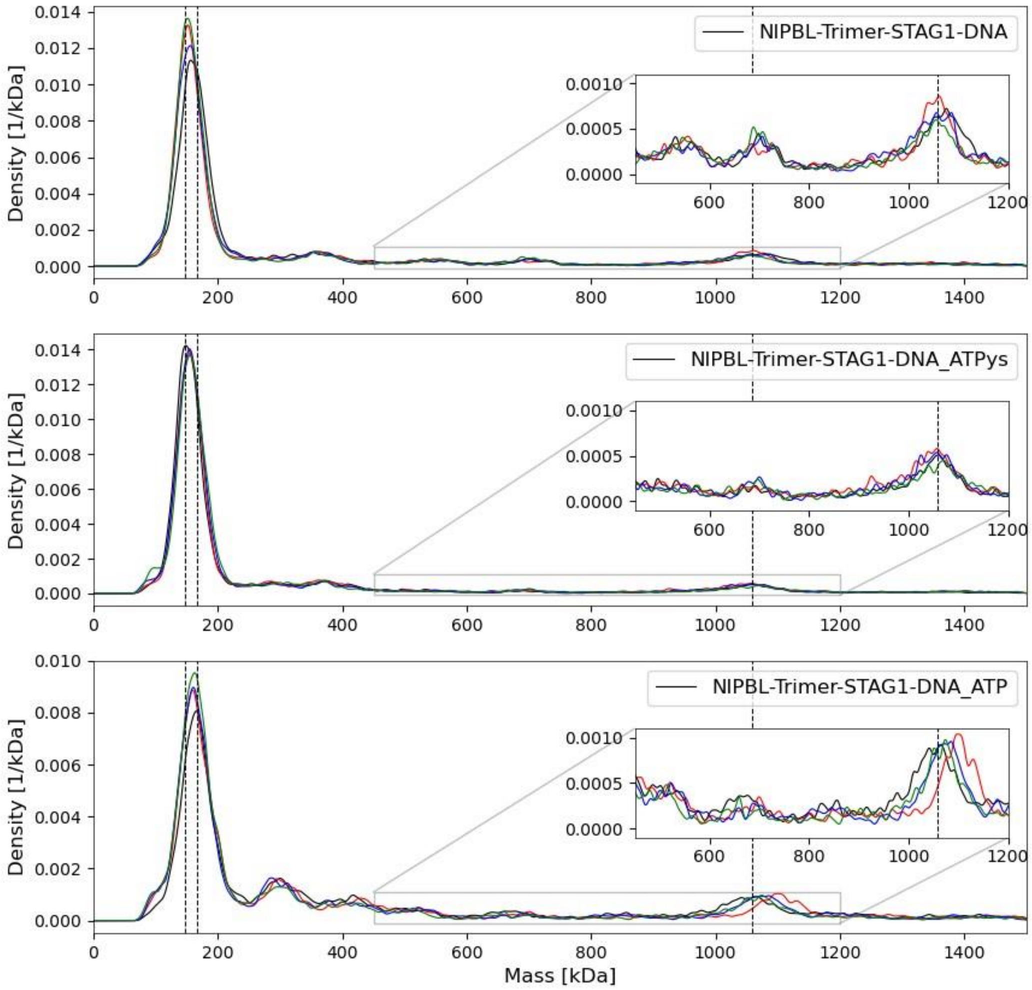


**Figure S17. Mass distributions for the STAG1-Trimer-NIPBL-DNA plots shown in Fig. 3E without subtraction.**

Unprocessed normalised experimental mass histograms of equimolar mixtures of NIPBL, Trimer, STAG1 and DNA following 5 min of incubation at room temperature in buffer containing 0 or 2.5 mM of ATP or ATPgS (as indicated in on the top right corner of each panel). The 3 technical repeats are shown with different colors. Vertical lines correspond to the expected masses of DNA and STAG1 that are forming the major peak and the expected mass of the complete complex. The average number of detected particles for the experiments in the 0-1500 kDa range for the technical repeats are $7.9 \times 10^3$ (APO, n = 3), $7.0 \times 10^3$ (ATPγs, n = 3), and $7.8\times 10^3$ (ATP, n = 3).

### Supplementary Tables

**Table S1. Mass fractions of the peaks of interests *vs.* the residuals.**

The table shows the results of dividing the mass histograms of the individual components (**Fig. S1**) into the peak of interest mass spectra and the residual spectra (see **Fig. S2**). The percentage indicates the relative fraction of detected molecules for each defined state. All spectra and fractions correspond to the detected mass between 0 and 1500 kDa.

| Component | Peak of Interest (%) | Residual (%) |
|---|---|---|
| MC302 | 75.9 | 24.1 |
| STAG1 | 91.6 | 8.4 |
| Trimer (batch 1) | 22.6 | 77.4 |
| Trimer (batch 2) | 41.3 | 58.7 |
| NIPBL | 49.2 | 50.8 |
| Truncated NIPBL | 69.3 | 30.7 |

**Table S2. Protein and DNA sequences**

| Protein | Sequence mass (kDa) | Sequence |
|---|---|---|
| SCC1(TEV)-Halo-His14 | 113 | MFYAHFVLSKRGPLAKIWLAAHWDKKLTKAHVFECNLESSVESIISPKVKMALRTSGHLLLGVVRIYHRK<br>AKYLLADCNEAFIKIKMAFRPGVVDLPEENREAAYNAITLPEEFHDFDQPLPDLDDIDVAQQFSLNQSRVE<br>EITMREEVGNISILQENDFGDFGMDDREIMREGSAFEDDDMLVSTTTSNLLLESEQSTSNLNEKINHLEYED<br>QYKDDNFGEGNDGGILDDKLISNNDGGIFDDPPALSEAGVMLPEQPAHDDMDEDDNVSMGGPDSPDSVD<br>PVEPMPTMTDQTTLVPNEEEAFALEPIDITVKETKAKRKRKLIVDSVKELDSKTIRAQLSDYSDIVTTLDLA<br>PPTKKLMMWKETGGVEKLFSLPAQPLWNNRLLKLFTRCLTPLVPEDLRKRRKGGEADNLDEFLKEFENPE<br>VPREDQQQQHQQRDVIDEPIIEEPSRLQESVMEASRTNIDESAMPPPENLYFQGPRENLYFQGASENLYFQG<br>PPQGVKRKAGQIDPEPVMPPQQVEQMEIPPVELPPEEPPNICQLIPELELLPEKEKEKEKEKEDDEEEEDEDA<br>SGGDQDQEERRWNKRTQQMLHGLQRALAKTGAESISLLELCRNTNRKQAAAKFYSFLVLKKQQAIELTQ<br>EEPYSDIIATPGPRFHIILEVLFQGPGGAGHMAEIGTGFPFDPHYVEVLGERMHYVDVGPRDGTPVLFLHGN<br>PTSSYVWRNIIPHVAPTHRCIAPDLIGMGKSDKPDLGYFFDDHVRFMDAFIEALGLEEVVLVIHDWGSALG<br>FHWAKRNPERVKGIAFMEFIRPIPTWDEWPEFARETFQAFRTTDVGRKLIIDQNVFIEGTLPMGVVRPLTEV<br>EMDHYREPFLNPVDREPLWRFPNELPIAGEPANIVALVEEYMDWLHQSPVPKLLFWGTPGVLIPPAEAARL<br>AKSLPNCKAVDIGPGLNLLQEDNPDLIGSEIARWLSTLEISGEPTTEDLYFSKHHHHSGHHHTGHHHHSGS<br>HHH |
| SMC1 | 143 | MGFLKLIEIENFKSYKGRQIIGPFQRFTAIIGPNGSGKSNLMDAISFVLGEKTSNLRVKTLRDLIHGAPVGKP<br>AANRAFVSMVYSEEGAEDRTFARVIVGGSSEYKINNKVVQLHEYSEELEKLGILIKARNFLVFQGAVESIA<br>MKNPKERTALFEEISRSGELAQEYDKRKKEMVKAEEDTQFNYHRKKNIAAERKEAKQEKEEADRYQRLK<br>DEVVRAQVQLQLFKLYHNEVEIEKLNKELASKNKEIEKDKKRMDKVEDELKEKKKELGKMMREQQQIEK<br>EIKEKDSELNQKRPQYIKAKENTSHKIKKLEAAKKSLQNAQKHYKKRKGDMDELEKEMLSVEKARQEFE<br>ERMEEESQSQGRDLTLEENQVKKYHRLKEEASKRAATLAQELEKFNRDQKADQDRLDLEERKKVETEAK<br>IKQKLREIEENQKRIEKLEEYITTSKQSLEEQKKLEGELTEEVEMAKRRIDEINKELNQVMEQLGDARIDRQ<br>ESSRQQRKAEIMESIKRLYPGSVYGRLIDLCQPTQKKYQIAVTKVLGKNMDAIIVDSEKTGRDCIQYIKEQR<br>GEPETFLPLDYLEVKPTDEKLRELKGAKLVIDVIRYEPPHIKKALQYACGNALVCDNVEDARRIAFGGHQR<br>HKTVALDGTLFQKSGVISGGASDLKAKARRWDEKAVDKLKEKKERLTEELKEQMKAKRKEAELRQVQS<br>QAHGLQMRLKYSQSDLEQTKTRHLALNLQEKSKLESELANFGPRINDIKRIIQSREREMKDLKEKMNQVE<br>DEVFEEFCREIGVRNIREFEEEKVKRQNEIAKKRLEFENQKTRLGIQLDFEKNQLKEDQDKVHMWEQTVK<br>KDENEIEKLKKEEQRHMKIIDETMAQLQDLKNQHLAKKSEVNDKNHEMEEIRKKLGGANKEMTHLQKEV<br>TAIETKLEQKRSDRHNLLQACKMQDIKLPLSKGTMDDISQEEGSSQGEDSVSGSQRISSIYAREALIEIDYGD<br>LCEDLKDAQAEEEIKQEMNTLQQKLNEQQSVLQRIAAPNMKAMEKLESVRDKFQETSDEFEAARKRAKK<br>AKQAFEQIKKERFDRFNACFESVATNIDEIYKALSRNSSAQAFLGPENPEEPYLDGINYNCVAPGKRFRPM<br>DNLSGGEKTVAALALLFAIHSYKPAPFFVLDEIDAALDNTNIGKVANYIKEQSTCNFQAIVISLKEEFYTKA<br>ESLIGVYPEQGDCVISKVLTFDLTKYPDANPNPNEQ |
| SMC3-Flag | 143 | MYIKQVIIQGFRSYRDQTIVDPFSSKHNVIVGRNGSGKSNFFYAIQFVLSDEFSHLRPEQRLALLHEGTGPRV<br>ISAFVEIIFDNSDNRLPIDKEEVSLRRVIGAKKDQYFLDKKMVTKNDVMNLLESAGFSRSNPYYIVKQGKIN<br>QMATAPDSQRLKLLREVAGTRVYDERKEESISLMKETEGKREKINELLKYIEERLHTLEEEKEELAQYQK<br>WDKMRRALEYTIYNQELNETRAKLDELSAKRETSGEKSRQLRDAQQDARDKMEDIERQVRELKTKISAM<br>KEEKEQLSAERQEQIKQRTKLELKAKDLQDELAGNSEQRKRLLKERQKLLEKIEEKQKELAETEPKFNSVK<br>EKEERGIARLAQATQERTDLYAKQGRGSQFTSKEERDKWIKKELKSLDQAINDKKRQIAAIHKDLEDTEA<br>NKEKNLEQYNKLDQDLNEVKARVEELDRKYYEVKNKKDELQSERNYLWREENAEQQALAAKREDLEK<br>KQQLLRAATGKAILNGIDSINKVLDHFRRKGINQHVQNGYHGIVMNNFECEPAFYTCVEVTAGNRLFYHI<br>VDSDEVSTKILMEFNKMNLPGEVTFLPLNKLDVRDTAYPETNDAIPMISKLRYNPRFDKAFKHVFGKTLIC<br>RSMEVSTQLARAFTMDCITLEGDQVSHRGALTGGYYDTRKSRLELQKDVRKAEEELGELEAKLNENLRR<br>NIERINNEIDQLMNQMQQIETQQRKFKASRDSILSEMKMLKEKRQQSEKTFMPKQRSLQSLEASLHAMEST<br>RESLKAELGTDLLSQLSLEDQKRVDALNDEIRQLQQENRQLLNERIKLEGIITRVETYLNENLRKRLDQVE<br>QELNELRETEGGTVLTATTSELEAINKRVKDTMARSEDLDNSIDKTEAGIKELQKSMERWKNMEKEHMD |

| | | |
|---|---|---|
| | | AINHDTKELEKMTNRQGMLLKKKEECMKKIRELGSLPQEAFEKYQTLSLKQLFRKLEQCNTELKKYSHVN<br>KKALDQFVNFSEQKEKLIKRQEELDRGYKSIMELMNVLELRKYEAIQLTFKQVSKNFSEVFQKLVPGGKA<br>TLVMKKGDVEGSQSQDEGEGSGESERGSGSQSSVPSVDQFTGVGIRVSFTGKQGEMREMQQLSGGQKSL<br>VALALIFAIQKCDPAPFYLFDEIDQALDAQHRKAVSDMIMELAVHAQFITTTFRPELLESADKFYGVKFRN<br>KVSHIDVITAEMAKDFVEDDTTHGSGGSDYKDDDDK |
| Flag-Halo-NIPBL-His | 353 | MDYKDDDDKSGGSMAEIGTGFPFDPHYVEVLGERMHYVDVGPRDGTPVLFLHGNPTSSYVWRNIIPHVA<br>PTHRCIAPDLIGMGKSDKPDLGYFFDDHVRFMDAFIEALGLEEVVLVIHDWGSALGFHWAKRNPERVKGI<br>AFMEFIRPIPTWDEWPEFARETFQAFRTTDVGRKLIIDQNVFIEGTLPMGVVRPLTEVEMDHYREPFLNPVD<br>REPLWRFPNELPIAGEPANIVALVEEYMDWLHQSPVPKLLFWGTPGVLIPPAEAARLAKSLPNCKAVDIGP<br>GLNLLQEDNPDLIGSEIARWLSTLEISGSGGSGSGSNGDMPHVPITTLAGIASLTDLLNQLPLPSPLPATTTKS<br>LLFNARIAEEVNCLLACRDDNLVSQLVHSLNQVSTDHIELKDNLGSDDPEGDIPVLLQAVLARSPNVFREK<br>SMQNRYVQSGMMMSQYKLSQNSMHSSPASSNYQQTTISHSPSSRFVPPQTSSGNRFMPQQNSPVPSPYAP<br>QSPAGYMPYSHPSSYTTHPQMQQASVSSPIVAGGLRNIHDNKVSGPLSGNSANHHADNPRHGSSEDYLHM<br>VHRLSSDDGDSSTMRNAASFPLRSPQPVCSPAGSEGTPKGSRPPLILQSQSLPCSSPRDVPPDILLDSPERKQ<br>KKQKKMKLGKDEKEQSEKAAMYDIISSPSKDSTKLTLRLSRVRSSDMDQQEDMISGVENSNVSENDIPFN<br>VQYPGQTSKTPITPQDINRPLNAAQCLSQQEQTAFLPANQVPVLQQNTSVAAKQPQTSVVQNQQQISQQGP<br>IYDEVELDALAEIERIERESAIERERFSKEVQDKDKPLKKRKQDSYPQEAGGATGGNRPASQETGSTGNGS<br>RPALMVSIDLHQAGRVDSQASITQDSDSIKKPEEIKQCNDAPVSVLQEDIVGSLKSTPENHPETPKKKSDPE<br>LSKSEMKQSESRLAESKPNENRLVETKSSENKLETKVETQTEELKQNESRTTECKQNESTIVEPKQNENRLS<br>DTKPNDNKQNNGRSETTKSRPETPKQKGESRPETPKQKSDGHPETPKQKGDGRPETPKQKGESRPETPKQ<br>KNEGRPETPKHRHDNRRDSGKPSTEKKPEVSKHKQDTKSDSPRLKSERAEALKQRPDGRSVSESLRRDHD<br>NKQKSDDRGESERHRGDQSRVRRPETLRSSSRNEHGIKSDSSKTDKLERKHRHESGDSRERPSSGEQKSRP<br>DSPRVKQGDSNKSRSDKLGFKSPTSKDDKRTEGNKSKVDTNKAHPDNKAEFPSYLLGGRSGALKNFVIPKI<br>KRDKDGNVTQETKKMEMKGEPKDKVEKIGLVEDLNKGAKPVVVLQKLSLDDVQKLIKDREDKSRSSLKP<br>IKNKPSKSNKGSIDQSVLKELPPELLAEIESTMPLCERVKMNKRKRSTVNEKPKYAEISSDEDNDSDEAFES<br>SRKRHKKDDDKAWEYEERDRRSSGDHRRSGHSHEGRRSSGGGRYRNRSPSDSDMEDYSPPPSLSEVARK<br>MKKKEKQKKRKAYEPKLTPEEMMDSSTFKRFTASIENILDNLEDMDFTAFGDDDEIPQELLLGKHQLNEL<br>GSESAKIKAMGIMDKLSTDKTVKVLNILEKNIQDGSKLSTLLNHNNDTEEEERLWRDLIMERVTKSADAC<br>LTTINIMTSPNMPKAVYIEDVIERVIQYTKFHLQNTLYPQYDPVYRLDPHGGGLLSSKAKRAKCSTHKQRV<br>IVMLYNKVCDIVSSLSELLEIQLLTDTTILQVSSMGITPFFVENVSELQLCAIKLVTAVFSRYEKHRQLILEEI<br>FTSLARLPTSKRSLRNFRLNSSDMDGEPMYIQMVTALVLQLIQCVVHLPSSEKDSNAEEDSNKKIDQDVVI<br>TNSYETAMRTAQNFLSIFLKKCGSKQGEEDYRPLFENFVQDLLSTVNKPEWPAAELLLSLLGRLLVHQFSN<br>KSTEMALRVASLDYLGTVAARLRKDAVTSKMDQGSIERILKQVSGGEDEIQQLQKALLDYLDENTETDPS<br>LVFSRKFYIAQWFRDTTLETEKAMKSQKDEESSEGTHHAKEIETTGQIMHRAENRKKFLRSIIKTTPSQFST<br>LKMNSDTVDYDDACLIVRYLASMRPFAQSFDIYLTQILRVLGENAIAVRTKAMKCLSEVVAVDPSILARLD<br>MQRGVHGRLMDNSTSVREAAVELLGRFVLCRPQLAEQYYDMLIERILDTGISVRKRVIKILRDICIEQPTFP<br>KITEMCVKMIRRVNDEEGIKKLVNETFQKLWFTPTPHNDKEAMTRKILNITDVVAACRDTGYDWFEQLLQ<br>NLLKSEEDSSYKPVKKACTQLVDNLVEHILKYEESLADSDNKGVNSGRLVACITTLFLFSKIRPQLMVKHA<br>MTMQPYLTTKCSTQNDFMVICNVAKILELVVPLMEHPSETFLATIEEDLMKLIIKYGMTVVQHCVSCLGA<br>VVNKVTQNFKFVWACFNRYYGAISKLKSQHQEDPNNTSLLTNKPALLRSLFTVGALCRHFDFDLEDFKGN<br>SKVNIKDKVLELLMYFTKHSDEEVQTKAIIGLGFAFIQHPSLMFEQEVKNLYNNILSDKNSSVNLKIQVLKN<br>LQTYLQEEDTRMQQADRDWKKVAKQEDLKEMGDVSSGMSSSIMQLYLKQVLEAFFHTQSSVRHFALNVI<br>ALTLNQGLIHPVQCVPYLIAMGTDPEPAMRNKADQQLVEIDKKYAGFIHMKAVAGMKMSYQVQQAINTC<br>LKDPVRGFRQDESSSALCSHLYSMIRGNRQHRRAFLISLLNLFDDTAKTDVTMLLYIADNLACFPYQTQEE<br>PLFIMHHIDITLSVSGSNLLQSFKESMVKDKRKERKSSPSKENESSDSEEEVSRPRKSRKRVDSDSDSDSEDD<br>INSVMKCLPENSAPLIEFANVSQGILLLLMLKQHLKNLCGFSDSKIQKYSPSESAKVYDKAINRKTGVHFHP<br>KQTLDFLRSDMANSKITEEVKRSIVKQYLDFKLLMEHLDPDEEEEEGEVSASTNARNKAITSLLGGGSPKN<br>NTAAETEDDESDGEDRGGGTSGSLRRSKRNSDSTELAAQMNESVDVMDVIAICCPKYKDRPQIARVVQKT<br>SSGFSVQWMAGSYSGSWTEAKRRDGRKLVPWVDTIKESDIIYKKIALTSANKLTNKVVQTLRSLYAAKDG<br>TSSSSGGSSHHHHHHHHHH |
| STAG1-His10 | 146 | MHHHHHHHHHHSGGSMITSELPVLQDSTNETTAHSDAGSELEETEVKGKRKRGRPGRPPSTNKKPRKSPG<br>EKSRIEAGIRGAGRGRANGHPQQNGEGEPVTLFEVVKLGKSAMQSVVDDWIESYKQDRDIALLDLINFFIQ<br>CSGCRGTVRIEMFRNMQNAEIIRKMTEEFDEDSGDYPLTMPGPQWKKFRSNFCEFIGVLIRQCQYSIIYDEY<br>MMDTVISLLTGLSDSQVRAFRHTSTLAAMKLMTALVNVALNLSIHQDNTQRQYEAERNKMIGKRANERL<br>ELLLQKRKELQENQDEIENMMNSIFKGIFVHRYRDAIAEIRAICIEEIGVWMKMYSDAFLNDSYLKYVGWT<br>LHDRQGEVRLKCLKALQSLYTNRELFPKLELFTNRFKDRIVSMTLDKEYDVAVEAIRLVTLILHGSEEALS<br>NEDCENVYHLVYSAHRPVAVAAGEFLHKKLFSRHDPQAEEALAKRRGRNSPNGNLIRMLVLFFLESELHE<br>HAAYLVDSLWESSQELLKDWECMTELLLEEPVQGEEAMSDRQESALIELMVCTIRQAAEAHPPVGRGTG<br>KRVLTAKERKTQIDDRNKLTEHFIITLPMLLSKYSADAEKVANLLQIPQYFDLEIYSTGRMEKHLDALLKQI<br>KFVVEKHVESDVLEACSKTYSILCSEEYTIQNRVDIARSQLIDEFVDRFNHSVEDLLQEGEEADDDDIYNVL<br>STLKRLTSFHNAHDLTKWDLFGNCYRLLKTGIEHGAMPEQIVVQALQCSHYSILWQLVKITDGSPSKEDLL<br>VLRKTVKSFLAVCQQCLSNVNTPVKEQAFMLLCDLLMIFSHQLMTGGREGLQPLVFNPDTGLQSELLSFV<br>MDHVFIDQDEENQSMEGDEEDEANKIEALHKRRNLLAAFSKLIIYDIVDMHAAADIFKHYMKYYNDYGDI<br>IKETLSKTRQIDKIQCAKTLILSLQQLFNELVQEQGPNLDRTSAHVSGIKELARRFALTFGLDQIKTREAVAT<br>LHKDGIEFAFKYQNQKGQEYPPPNLAFLEVLSEFSSKLLRQDKKTVHSYLEKFLTEQMMERREDVWLPLIS<br>YRNSLVTGGEDDRMSVNSGSSSSKTSSVRNKKGRPPLHKKRVEDESLDNTWLNRTDTMIQTPGPLPAPQL<br>TSTVLRENSRPMGDQIQEPESEHGSEPDFLHNPQMQISWLGQPKLEDLNRKDRTGMNYMKVRTGVRHAV<br>RGLMEEDAEPIFEDVMMSSRSQLEDMNEEFEDTMVIDLPPSRNRRERAELRPDFFDSAAIIEDDSGFGMPM<br>F |
| TwinStrep-Flag-Halo-ΔN- | 243 | MSAWSHPQFEKGGGSGGGSGGSAWSHPQFEKSGGSDYKDDDDKSSGGSEIGTGFPFDPHYVEVLGERMH<br>YVDVGPRDGTPVLFLHGNPTSSYVWRNIIPHVAPTHRCIAPDLIGMGKSDKPDLGYFFDDHVRFMDAFIEA<br>LGLEEVVLVIHDWGSALGFHWAKRNPERVKGIAFMEFIRPIPTWDEWPEFARETFQAFRTTDVGRKLIIDQ<br>NVFIEGTLPMGVVRPLTEVEMDHYREPFLNPVDREPLWRFPNELPIAGEPANIVALVEEYMDWLHQSPVP<br>KLLFWGTPGVLIPPAEAARLAKSLPNCKAVDIGPGLNLLQEDNPDLIGSEIARWLSTLEISGSRNGDMPHVP<br>ITTLAGIASLTDGSIDQSVLKELPPELLAEIESTMPLCERVKMNKRKRSTVNEKPKYAEISSDEDNDSDEAFE |

| | | |
|---|---|---|
| NIPBL-His10 | | SSRKRHKKDDDKAWEYEERDRRSSGDHRRSGHSHEGRRSSGGGRYRNRSPSDSDMEDYSPPPSLSEVARK<br>MKKKEKQKKRKAYEPKLTPEEMMDSSTFKRFTASIENILDNLEDMDFTAFGDDDEIPQELLLGKHQLNEL<br>GSESAKIKAMGIMDKLSTDKTVKVLNILEKNIQDGSKLSTLLNHNNDTEEEERLWRDLIMERVTKSADAC<br>LTTINIMTSPNMPKAVYIEDVIERVIQYTKFHLQNTLYPQYDPVYRLDPHGGGLLSSKAKRAKCSTHKQRV<br>IVMLYNKVCDIVSSLSELLEIQLLTDTTILQVSSMGITPFFVENVSELQLCAIKLVTAVFSRYEKHRQLILEEI<br>FTSLARLPTSKRSLRNFRLNSSDMDGEPMYIQMVTALVLQLIQCVVHLPSSEKDSNAEEDSNKKIDQDVVI<br>TNSYETAMRTAQNFLSIFLKKCGSKQGEEDYRPLFENFVQDLLSTVNKPEWPAAELLLSLLGRLLVHQFSN<br>KSTEMALRVASLDYLGTVAARLRKDAVTSKMDQGSIERILKQVSGGEDEIQQLQKALLDYLDENTETDPS<br>LVFSRKFYIAQWFRDTTLETEKAMKSQKDEESSEGTHHAKEIETTGQIMHRAENRKKFLRSIIKTTPSQFST<br>LKMNSDTVDYDDACLIVRYLASMRPFAQSFDIYLTQILRVLGENAIAVRTKAMKCLSEVVAVDPSILARLD<br>MQRGVHGRLMDNSTSVREAAVELLGRFVLCRPQLAEQYYDMLIERILDTGISVRKRVIKILRDICIEQPTFP<br>KITEMCVKMIRRVNDEEGIKKLVNETFQKLWFTPTPHNDKEAMTRKILNITDVVAACRDTGYDWFEQLLQ<br>NLLKSEEDSSYKPVKKACTQLVDNLVEHILKYEESLADSDNKGVNSGRLVACITTLFLFSKIRPQLMVKHA<br>MTMQPYLTTKCSTQNDFMVICNVAKILELVVPLMEHPSETFLATIEEDLMKLIIKYGMTVVQHCVSCLGA<br>VVNKVTQNFKFVWACFNRYYGAISKLKSQHQEDPNNTSLLTNKPALLRSLFTVGALCRHFDFDLEDFKGN<br>SKVNIKDKVLELLMYFTKHSDEEVQTKAIIGLGFAFIQHPSLMFEQEVKNLYNNILSDKNSSVNLKIQVLKN<br>LQTYLQEEDTRMQQADRDWKKVAKQEDLKEMGDVSSGMSSSIMQLYLKQVLEAFFHTQSSVRHFALNVI<br>ALTLNQGLIHPVQCVPYLIAMGTDPEPAMRNKADQQLVEIDKKYAGFIHMKAVAGMKMSYQVQQAINTC<br>LKDPVRGFRQDESSSALCSHLYSMIRGNRQHRRAFLISLLNLFDDTAKTDVTMLLYIADNLACFPYQTQEE<br>PLFIMHHIDITLSVSGSNLLQSFKESMVKDKRKERKSSPSKENESSDSEEEVSRPRKSRKRVDSDSDSDSEDD<br>INSVMKCLPENSAPLIEFANVSQGILLLLMLKQHLKNLCGFSDSKIQKYSPSESAKVYDKAINRKTGVHFHP<br>KQTLDFLRSDMANSKITEEVKRSIVKQYLDFKLLMEHLDPDEEEEEGEVSASTNARNKAITSLLGGGSPKN<br>NTAAETEDDESDGEDRGGGTSGSLRRSKRNSDSTELAAQMNESVDVMDVIAICCPKYKDRPQIARVVQKT<br>SSGFSVQWMAGSYSGSWTEAKRRDGRKLVPWVDTIKESDIIYKKIALTSANKLTNKVVQTLRSLYAAKDG<br>TSSSGGSHHHHHHHHHH |
| MC302 | 186 kDa | CATATGTGCAGTCCCCCCATAAAAAAACCCGCCGAAGCGGGTTTTTACGTTATTTGCGGATTTTAATT<br>AACTAGTTCTAGAGCTCACGTGCAGCTGTCGACGGTACCTGCAGGCCGGCCAGTACTGGGCCCCAAC<br>AGATAAAACGAAAGGCCCAGTCTTTCGACTGAGCCTTTCGTTTTATTGGGCCCCCTTGGTCAAATTGG<br>GTATACCCATTTGGGCCTGGTCTGGCCGGCATGGCGCATTACAGCGGTACGCAATTTAAATGCGCCTA<br>GCGCATTTTCCCGACCTTAATGCGCCTCGCG |
